# Chromosomal rearrangements with stable repertoires of genes and transposable elements in an invasive forest-pathogenic fungus

**DOI:** 10.1101/2021.03.09.434572

**Authors:** Arthur Demené, Benoît Laurent, Sandrine Cros-Arteil, Christophe Boury, Cyril Dutech

## Abstract

Chromosomal rearrangements have been largely described among eukaryotes, and may have important consequences on evolution of species. High genome plasticity has been often reported in Fungi, which may explain their apparent ability to quickly adapt to new environments. *Cryphonectria parasitica*, causing the Chestnut blight disease, is an invasive fungal pathogen species associated with several recent host shifts during its successive introductions from Asia to North America and Europe. Previous cytological karyotyping and genomic studies suggested several chromosomal rearrangements which remains to be described in detail for this species. A serious limitation for valid genome comparisons is the access to robust genome assemblies that usually contain genomic regions of low complexity. We present a new de novo whole-genome assembly obtained from a new method of DNA extraction and long-reads sequencing Nanopore technology obtained from a Japanese isolate sampled in the native area of the species. The comparison with a recently published reference genome revealed stable gene and transposable elements (TEs) repertoires. We also showed that the *C. parasitica* genome is lowly compartmentalized, with a poor association between TEs and genes, such as those potentially involved in host interactions (i.e., genes coding for small secreted proteins or for secondary metabolites). This genome comparison, however, detected several large chromosomal rearrangements that may have important consequences in gene regulations and sexual mating in this invasive species. This study opens the way for more comparisons of high-quality assembled genomes, and questions the role of structural variations in the invasive success of this fungal pathogen species.

## Introduction

Large structural variations of genome (i.e., translocation, insertion, deletions and inversions greater than 100kb) are more and more described in numerous organisms and may have important consequences on species adaptation (Alonge et al. 2020, Wellenreuther et al. 2019). Filamentous microorganisms (i.e., fungi and oomycetes) have generally small genome sizes, present a large diversity of lifestyles, and several of them can be easy to manipulate in laboratory to characterize their phenotype. These general characteristics make them especially interesting to study genomic evolution in eukaryotes and molecular mechanisms involved in genetic adaptation (Raffaele and Kamoun 2012, Chang et al. 2013, Wang et al. 2020, Yadav et al. 2020). For example, fungal genomic studies have investigated gene repertory associated with specific lifestyles (e.g. Spanu et al. 2010, Duplessis et al. 2011, Riley et al. 2014), evolution of sexual chromosomes (Badouin et al. 2015), signatures of selection along the genome (Badouin et al. 2017), recombination hot-spots (Croll et al. 2015, Laurent et al. 2018), speciation or host specialization (e.g. Stukenbrock and Bataillon 2012, Hou et al. 2014), and cryptic genetic exchanges among genetic lineages (Milgroom et al. 2014, Demené et al. 2019) or between species (Cheeseman et al. 2014, Dhillon et al. 2015). One of the major findings of these studies was the high genomic plasticity observed within and between closely related species, associated with various genetic mechanisms, such as intra- and inter-chromosomal rearrangements (Hou et al. 2014, Frantzeskakis et al. 2018, Plissonneau et al. 2016, Tsushima et al. 2019), gains or losses of genes (Yoshida et al. 2016, Hartmann et al. 2017), variation in chromosome numbers (Ma et al. 2010), or chromosome stretching (Grandaubert et al. 2014). Like single nucleotide polymorphisms, many of these genomic structural variations have been considered as fuel for genetic adaptation of fungal species (Raffaele and Kamoun 2012).

These studies have also lead to the development of the « two-speed genome » concept (Grandaubert et al. 2014, Dong et al. 2015) also related to the “genomic island” concept (Fedorova et al. 2008), “genome compartmentalization” (Dutheil et al. 2016, Frantzeskakis et al. 2019), or genomes with “dynamic compartments” (Torres et al. 2020). The general idea behind these concepts is that the genomes of many fungal pathogens are composed of two distinct compartments. One compartment, often enriched in transposable elements (TE) and poor in genes, presents a high rate of structural variations, copy-number variations, genetic recombination, sequence polymorphisms or genes under strong positive selection (Wang et al. 2017, Faino et al. 2016, Laurent et al. 2018, Sánchez-Vallet et al. 2018, Eschenbrenner et al. 2020, Torres et al. 2020). A second, rich in genes and poor in TEs, is considered less dynamic. Genes in close physical proximity to these TEs have often been described as genes encoding proteins potentially involved in plant pathogen interactions (Raffaele and Kamoun 2012). Their proximity with TEs has been considered as an advantage for rapid diversification and adaptation to new hosts species or genotypes, because of direct (gene or promoter disruption, introduction of regulatory information) and indirect (induced mutations, ectopic rearrangement mediation, or heterochromatin formation) effects of TEs on neighbour genes (Daboussi and Capy 2003, Seidl and Thomma 2017). Consequently, TEs may mediate gain or loss of genes involved in virulence against host (e.g. Castanera et al. 2016, Frantzeskakis et al. 2019). However, this hypothesis has been mainly proposed for fungal pathogens infecting crops for which a need of rapid adaption is often assumed (Brown and Tellier 2011, Raffaele and Kamoun 2012). There is less evidence that this genomic architecture, as well as important variations in gene contents, may be observed in fungal plant pathogens living in more natural ecosystems, and for which selective pressures probably differ (see for example Hartmann et al. 2018). Therefore, the role of TEs in genomic adaptation remains largely debated, certainly because of the scarcity of studies comparing multiple genomes at different evolutionary timescales and lifestyles (Muszewska et al. 2019). As a matter of fact, comparisons of whole-genomes sequences have been rather limited until recently, especially due to the difficulties to assemble regions of low complexity (including TE clusters; e.g. Badouin et al. 2015, Frantzeskakis et al. 2018). The recent emergence of the third generation of sequencing technology (e.g. Oxford Nanopore Technologies or Pacific Biosciences sequencing platforms), yielding sequences of several kilobases, allows to solve these limitations (Badouin et al. 2015, Plissonneau et al. 2018). These technological advances are becoming much more accessible for large numbers of taxa, opening new perspectives to investigate the extend and the molecular mechanisms at the origin of genomic structural variations, as well as their phenotypic effects in different ecological and evolutionary contexts (Alonge et al. 2020, Badet et al. 2020).

*Cryphonectria parasitica* is a devastating forest fungal pathogen infecting chestnut species (*Castanea* sp.) over the world (Anagnostakis 1987). The species originating from Asia has been introduced several times in North America and Europe where it caused serious outbreaks and resulted in the disappearance of the mature *C. dentata* in North America (Milgroom et al. 1996, Dutech et al. 2012). The invasive success observed in the two introduced areas, as well as the associated host shifts (*C. crenata* and *C. molissima* in Asia, *C. dentata* in North America and *C. sativa* in Europe, or oaks (Quercus sp.) in different countries), asks the question of the adaptative processes associated to these events. For example, change in reproductive mode, a common feature of fungal invasions (Gladieux et al. 2015), has been observed between native and introduced areas (Milgroom et al. 2008, Dutech et al. 2010). It is not clear whether this reproductive shift is a cause or a consequence of invasive success, and due to genetic incompatibilities among the introduced genetic lineages (Dutech et al. 2010). Recently, a high-quality reference genome of *C. parasitica* was published from the laboratory reference strain EP155, originally sampled in North-American from *C. dentata* (Crouch et al. 2020). This reference genome offers the opportunity to study genomic changes that may have occurred during the different introductions of *C. parasitica*. We previously noted that this first reference genome was, in some degrees, different from a new de novo genome assembly obtained from a French isolate (YVO003), using a long-reads sequencing technology (Demené et al. 2019). This result was consistent with a previous cytological karyotyping study comparing the strain of the reference genome (EP155) with other strains sampled in North America, that suggested several chromosomal rearrangements (Eusebio-Cope et al. 2009). Overall, these results suggest that *C. parasitica* genomes can be quite different, and these differences could be linked to its invasion history. However, the de novo genome assembly obtained on YVO003 was slightly shorter than the EP155 reference genome, suggesting that some parts of the genome were missing, and precluding an accurate comparison of the two genome assemblies.

To study putative chromosomal rearrangements and other structural genomic variations which may have occurred between native and introduced *C. parasitica* genomes, we sequenced a whole-genome of a *C. parasitica* strain, sampled by Liu et al. (2003) in the native Japanese area, and compared the obtained de novo assembly to the EP155 North-American reference genome (Crouch et al. 2020). The new genome sequence of this strain was produced using a strategy of different sequencing technologies, mixing the use of Oxford Nanopore Technology (ONT) and Illumina sequencing, in order to obtain a robust assembly of regions of low complexity. DNA quality being crucial in the success of ONT sequencing, we developed a new protocol for extracting high molecular weight DNA. Using such a strategy, we produced a near-chromosome level assembly, which have been manually curated, of the Japanese ESM015 *C. parasitica* isolate. We compared this assembly with the recently published reference EP155 genome (Crouch et al. 2020) to estimate the importance of large structural variations between the two genomes. We produced a gene prediction on the new de novo assembly, and a manually curated set of TEs representative of both genome assemblies. We looked for possible changes in composition or TEs and gene repertoires which may have occurred during the recent host shift between the Asian *Castanea* species and the North American *C. dentata* species, and may have contributed to the invasive success of the species.

## Methods

### Sequenced isolate, extraction DNA protocol and sequencing

The sequenced strain ES15 (called ESM015 in this study and in Demené et al. 2019) was sampled in the Japanese Esashi population in 1998 on *C. crenata*, and isolated from a single asexual spore (Liu et al. 2003). For this sequencing, this monosporous isolate was multiplied again on potato dextrose agar culture as described in Hoegger et al. (2000). To obtain high quality DNA required for long-run sequencing, a new protocol of DNA extraction was developed that aimed to deal with important contents in polysaccharides present in many fungal species (Möller et al. 1992). Briefly, a six-day-old cultivated mycelium (total weight: ~0.4g) was grinded using liquid nitrogen and pre-chilled mortar and pestle. Extraction solution consisted of 10 % Sodium Dodecyl Sulfate lysis buffer with 2 mg of proteinase K at 20 mg.mL-1. Grinded mycelium was homogenized with the extraction buffer. No vortexing was applied for maximizing the chance of obtaining a high molecular weight DNA. Two steps of phenol chloroform isoamyl alcohol 25:24:1 were performed, with an addition of RNAse A for 2% of the final volume. Four ethanol (70%) washes of the DNA pellet have been carefully made to avoid breaking DNA. In order to remove the remaining polysaccharides, a final purification step was performed: the DNA solution was brought to 1.2M NaCl solution, and polysaccharides were precipitated and isolated from DNA using a diethyl ether solution saturated with DNase-free water. The complete protocol is available in the supplementary data (Sup. Mat. 1, also available on dx.doi.org/10.17504/protocols.io.bzbdp2i6). DNA purification was performed using 1.0X magnetic beads (11% polyethylen glycol), followed by DNA size selection (>1kb) and cleaning, using an optimized solid-phase reversible immobilization according to Fisher et al. (2011), and a bead mixture according to Schalamun et al. (2019). Then, 16μg of purified gDNA was sheared to approximately 40Kb using hydropore-long and megaruptor II (Diagenode, Seraing, Belgium), end-repaired, purified and size selected over 20Kb by using four lanes of a High-pass plus 0.75% agarose-gel cassette and blue pippin instrument (Sage Science, Beverly, MA, USA) and then again purified. A 1D library preparation was performed according to Oxford Nanopore technology recommendations using SQK-LSK109 sequencing kit. Sequencing was performed with a R9.4.1 Flowcell on the GridION device using MinKNOW (version 2.2), and a dedicated basecaller guppy GridION version 1.8.5.1. (Oxford Nanopore technology, Cambridge, UK), at the Genome Transcriptome Facility of Bordeaux (PGTB, University of Bordeaux, France). For increasing the quality of the genome assembly, Illumina reads of the ESM015 isolate produced in Demené et al. (2019) were added to the analysis. These reads were trimmed and filtered using prinseq v0.20.4 (Schmieder and Edwards 2011). First, the fifth first 5’ nucleotides were trimmed due to skewed base composition introduced by sequencing preparation, and duplicate reads were removed. Reads with an overall mean Phred-scaled value less than 30 were discarded. Remaining reads were further 3’ and 5’ trimmed for quality (Phred-scaled threshold of 20). Finally, reads with size lower than 50bp or exceeding 280 bp were excluded.

### Comparison of genome assemblers

To produce a high confidence de novo assembly genome, several pipelines based on different strategies were tested: 1) Assembly of short reads previously produced for this monosporous isolate (Demené et al. 2019), followed by the mapping of long reads to close gaps of the assembly graph (HybridSPAdes using the De-Bruijn graph (DBG) algorythm; Antipov et al. 2016), 2) Assembly of long reads corrected using short reads (Ra using the Overlap Layout Consensus (OLC) algorythm; https://github.com/lbcb-sci/ra), or 3) Assembly of raw long reads followed by a polishing of the assembly using the short reads (Miniasm using the OLC algorythm; Li 2016). For HybriSPAdes, an optimized k-mer size has to be determined to close gaps, depending on the read length and depth. We tested six different K-mer values (33, 55, 77, 99, 111 and 127) and used Bandage (https://github.com/rrwick/Bandage/wiki/Effect-of-kmer-size) to identify the best k-mer value (Marijon et al. 2019). We choose the k-mer size producing the assembly graph with the lowest complexity (lowest number of nodes and edges and highest nodes N50), while checking that the graph was not broken into too much disconnected parts as suggested on the Bandage user’s manual (https://github.com/rrwick/Bandage/wiki/Effect-of-kmer-size). HybridSPAdes was ran on R1 and R2 paired-end reads corrected with the short read error correction step (BayesHammer; Nikolenko et al. 2013), the accurate mode on, and the Mismatch correction step. The Ra assembler was run on the complete ONT raw data, with the corrected unpaired R1 Illumina reads (having a higher quality than R2), and with basic parameters and option -x ont. For Miniasm, highly sensitive to sequencing errors, an hybrid correction step of the raw Nanopore long-reads with CoLoRMap (Haghshenas et al. 2016) has been added prior to runs. Miniasm was run on the complete set of corrected ONT reads with the parameters -ax ava-ont.

For each method, the following metrics were extracted to assess the quality of assemblies: N50, L50, genome size, total number of contigs, length of the largest contig, GC content. Completeness of genome assemblies was assessed by three methods: 1-searching for the 1,315 genes from the Ascomycota database using BUSCO (Simão et al. 2015), 2-performing a KAT plot for each assembly and estimating assembly completeness using the Kmer Analysis Toolkit (KAT) “comp” tool (https://kat.readthedocs.io/en/latest/index.html; Mapleson et al. 2017), and 3-mapping on each assembly the corrected paired-end reads Illumina libraries using Bowtie2 (Langmead and Salzberg 2012) with basic parameters.

### Manual curation of the new de novo genome assembly

A manual curation step was done on the assembly yielding the best assembly metrics. First, a scaffolding of contigs was performed using npScarf (Cao et al. 2017), and small contigs (i.e. < 1000bp) were removed. Second, long reads were then mapped on the scaffolds using minimap2 (Li 2018), as well as short reads using bowtie2 (Langmead and Salzberg 2012). Reads with a map quality of zero (i.e. mapped with an equal quality on at least two locations) were removed. Read depth (bamCoverage from Deeptools, Ramirez et al. 2016) and read quality (Samtools, Li et al. 2009) were then estimated to detect regions with less than two mapped long reads, long reads coverage higher than twice the mean coverage of the genome, and more than five mis-paired short reads. All these regions were manually checked using the Integrative Genomics Viewer (IGV, Thorvaldsdóttir et al. 2013), and manually cut when misassemblies were detected (i.e. regions with nonoverlapping long reads, mis-pairing of Illumina reads, or with outliers for assembly metrics). When these putative chimeric regions were associated with mis-paired short-reads, the pairing information was used to reassemble the cut parts and to construct new scaffolds. Finally, we looked at each end of the scaffolds for mis-paired reads to perform an additional manual scaffolding using these information. Three cycles of scaffolding with this manual curation were performed to obtain the final reference genome assembly. Finally, we used BUSCO v3 (Simão et al. 2015) with the Ascomycota odb9 single-copy orthologs set to re-assess genome assembly completeness. We mapped Illumina and Nanopore reads using Bowtie2 and minimap2 (see details in scripts). Coverage has been assessed using bamCoverage from Deeptools (Ramirez et al. 2016) with a RPKM normalization (Reads Per Kilobase per Million mapped reads) and a read filtering for a minimum mapping quality of 10. Telomeric regions were identified based on -TTAGGG- repeated sequences described for fungi (Casas-Vila et al. 2015), and of -TTAGGGCTAGGG- and -CTAGGG- known to be specifically associated to such regions in *C. parasitica* genome (Qi et al. 2013). Synteny between this assembly and the EP155 reference genome was assessed using nucmer (from MUMmer4, Marçais et al. 2018). Visualisation of the synteny was done using Circa (http://omgenomics.com/). Filtering of syntenic links > 10kb or with an identity < 95% has been made using the delta-filter program from MUMmer4.

### Prediction, annotation and curation of transposable elements

For the TEs detection, the REPET pipeline (https://urgi.versailles.inra.fr/Tools/REPET) was used on the curated assembly, including Tedenovo for the consensus prediction (Flutre et al. 2011, Hoede et al. 2014) and TEannot for their annotation (Quesneville et al. 2005). A manual curation step was performed on the TEs consensus library following the instructions of the REPET pipeline. The percentage of genome coverage by TEs were compared between the two TEs libraries (i.e. the libraries obtained before and after curation) in order to check whether no information has been lost. Transposable elements sequences without at least one full length copy (FLC) found in the genome were considered doubtfull, and were removed from the libraries. These sequences were considered as duplications, and likely related to FLC consensus sequences obtained during annotation. Part of the TE sequences which have been inferred from only one high-scoring segment pair (HSP) were cut. Each TE identification was validated according to its repeated extremities and genes content using the Wicker’s classification (Wicker et al. 2007). Two TE databases were created: TEs_db1 for ESM015, the curated one, and TEs_db2 for EP155. A final TEs data base (TEs_db3) was then created by gathering the two first data bases. This data base was composed of the complete Tes_db1 plus the TEs of Tes_db2, and covering at least 0.05% of one of the two genomes (i.e. 22kb). TEs_db3 was then used to re-annotate both genomes. Details of the curation of the TE annotation are given in Sup. Mat. 2. Divergence among pairs of TEs sequences annotated in the ESM015 genome was estimated by aligning them to each consensus using nucmer (Marçais et al. 2018).

In order to test whether a signal of repeat induced point (RIP) mutation, an active defense mechanism against the TE replication in fungi (Selker 1990), was present in the genome of *C. parasitica*, we first looked for the presence of AT-rich isochores in the ESM015 isolate genome, a typical feature of RIP-containing genomes (Gout et al. 2006). Second, we calculated two ratios of dinucleotides : 1) the “pre-RIP” ratio being the expected nucleotide ratio before the RIP mutation, and 2) the “post-RIP” ratio being the expected ratio of the product of the RIP mutation (i.e. (CpA+TpG)/(ApC+GpT) and TpA/ApT respectively; Margolin et al. 1998, Selker 1990). Transposable element sequences with a pre-RIP index lower than the baseline estimated from the genomic sequences masked for TEs, and a post-RIP index higher than the baseline, indicate a bias in AT frequencies in agreement with a RIP mutation activity (Margolin et al. 1998, Selker 1990). We also estimated a third index (CpG/GpC) specifically designed for *C. parasitica*, and measuring the depletion of the RIP on the basis of the preferential nucleotide targets in this species (see for details Clutterbuck 2011). Confidence intervals at 99.9% of the three baseline RIP indexes was estimated using a sliding window of 10kb over both masked genomes.

### Genes, effectors, small-secreted proteins (SSPs) and secondary metabolites (SM) annotation, and comparison of the two genomes

Gene prediction was performed on the curated genome using BRAKER v2.1.2 pipeline (Hoff et al. 2019). This pipeline first trained AUGUSTUS software (Stanke et al. 2008) using the gene Catalog 20091217 of the reference genome EP155 (Crouch et al. 2020). Three steps with default parameters were performed: 1) EP155 proteins aligned on the ESM015 genome assembly using GenomeThreader (Gremme et al. 2013), 2) Augustus training from the alignment file using NCBI BLAST (Altschul et al. 1990), and 3) Augustus gene prediction on the curated genome. Gene functions were predicted by i) searching for protein signatures on the different InterPro databases using Interproscan V5.29-68.0 (Jones et al. 2014) in cluster mode with standard parameters and ii) by comparing protein sequences with proteins described in the manually annotated SwissProt database (assessed the 18th of November 2020) using blastp (ncbi-blast-2.10.0+). Blastp results were mapped against the Gene Ontology Annotation Database using Blast2GO V5 (Götz et al. 2008). Annotations from InterproScan and BlastP were merged and annotated using Blast2Go V5. Secreted proteins were more specifically predicted using the deep neural network-based software SignalP version 5.0 (Armenteros et al. 2019). In this study, only the predictions with a likelihood of 0.5 or greater were considered as secreted proteins. In addition, genes encoding potential effectors (i.e. proteins or small molecules that are secreted from plant-associated organisms to the host and alter host-cell structure and function; Kamoun 2006) were predicted using all the secreted proteins predicted from SignalP (i.e. also including the proteins with a likelihood smaller than 0.5), and the machine learning based software EffectorP version 2.0 (Sperschneider et al. 2018). Only the effectors predicted with a likelihood of 0.6 or higher were considered. A set of candidate secreted effector proteins (CSEP) was also proposed by selecting the effectors from the set of the predicted secreted proteins having a likelihood greater than 0.5. Secondary metabolite biosynthesis gene clusters were identified and annotated using antiSMASH fungal version web tool (antiSMASH 5.0, Blin et al. 2019).

The predicted proteins content between the previously annotated reference genome (Crouch et al. 2020) and this new reference genome have been compared using OrthoFinder v2.4.0 (Emms and Kelly 2019) using default parameters. Differences in gene content were corrected for technical biases, mainly due to predictions using different tools in Crouch et al. (2020) and this study. The nuclear sequences of the predicted proteins showing no homologue in the other genome were searched on the other genome sequence using blastn (v2.9.0+). Using a conservative filter on these blast results, gene sequence alignments of the request were considered as non-functional either when starting after the start codon, aligning less than 80% in size with the target sequences, or showing alignments with nucleotide similarities < 80%.

## Results

### Comparison of the obtained assemblies for the ESM015 genome

Two pooled DNA extractions yield a total of 66.0 μg of double-stranded DNA (estimated in 110 μL of final solution with Qbit). The DNA quality, estimated with the ND8000 nanodrop, was respectively 1.87 and 1.45 for 260/280 and 260/230 ratio. The Nanopore sequencing produced 453,365 reads having a mean and median length of 9.40kb and 8.72kb respectively, N50 of 15.46kb, a mean quality of 8.0 (median 8.9), and a cumulative length of 4.26Gb (details on Sup. Mat. 3). Based on a 45Mb genome (Eusebio-Cope et al. 2009, Crouch et al. 2020), the estimated coverage of the genome is 94.7X. The Miniasm assembler using Nanopore reads led to an assembly size of 46.6Mb (N50 = 5.3Mb; L50 = 4), but the recovery rate for the 1,315 genes using BUSCO was only 0.5%, only half (53.15%) of Illumina reads properly mapped on this assembly (Table S1), and the KAT estimated assembly completeness was only 20.75% (All KAT plots for different assemblers and curation steps are in Fig. S1). Adding a preliminary correction step of long reads using paired-end Illumina reads with CoLoRMap significantly improved the results with a genome size (43.6Mb) close to the EP155 genome (43.8Mb), a better completeness for gene contents (BUSCO genes recovery rate of 69.2%, Table S1), and a better KAT assembly completeness estimate (72.41%). Using Ra, we obtained the shortest genome assembly (42.4Mb; N50 = 2.8Mb; L50 = 6), with a higher GC rate than in EP155 (51.3% and 50.8%, respectively). The completeness estimates of this genome was high (98.6% of the BUSCO genes recovered, 94.61% Illumina reads mapped, and a KAT assembly completeness estimate of 99.86%), but the de novo assembly was more fragmented than the assembly obtained with Miniasm (Table S1). For HybriSPAdes, we finally selected 127 as the optimized kmer size (see Table S2 for details). Although this assembly was the most fragmented of all those produced in this study (i.e. 235 contigs, maximum length 3.2Mb and minimum length 129bp), it yielded good results considering together the basic genome statistics (genome size of 43.4Mb close to the EP155 assembly, N50 = 1.9Mb, L50 = 9, and GC rate similar to EP155; Table S1), BUSCO analysis (recovering 98.7% of the genes of the Ascomycota database), the alignment rate of Illumina reads (96.68%) (Table S1), and the estimated assembly completeness using KAT (99.92%). Although close to the statistics obtained with RA, this last genome assembly was chosen for the following analyses.

### Detection and correction of several chimeric scaffolds through a manual curation

The scaffolding step using npScarf with long reads on the hybridSPAdes assembly allowed to obtain a new assembly with 113 scaffolds for a total length of 43.46Mb (L50=4, N50=3.66Mb). Ninety-three small scaffolds (< 10,000bp) were removed (~ 150kb total length, 47.7 % GC). Twenty chimeric regions were cut, and 15 of which have been anchored to the corresponding part using paired-end reads information. Eight new bonds were also determined between ends of the scaffolds using the paired-ends information. We identified the 148kb mitochondrial genome on the basis of a high coverage of ~ 1000X with Nanopore reads, and ~4000X with Illumina paired-end reads. On this assumed mitochondrial contig, Mitos2 (Donath et al. 2019) identified several genes homologues to mitochondrial genes, such as cytochrome b and c oxydase, NADH dehydrogenase, ATP synthase and RNA polymerase (see Table S3 for details). The final assembly after two others similar steps of scaffolding and manual curation yielded 17 scaffolds (plus the mitochondrial genome) for a total length of 43.31Mb (L50=5, N50=4,1Mb, Figure 1, Table 1). After all the curation steps, the assembly completeness estimated using KAT remained high (99.88%). The Nanopore long reads were mapped again on this final assembly, and no break of mapping for these long reads was detected (Figure 1). The assembled scaffolds range from 39kb (ESM_16) to 5.38Mb (ESM_2). Fourteen telomeric regions were found at the end of ten scaffolds, with four scaffolds (ESM_2, ESM_3, ESM_4 and ESM_6) having telomeric motives on both ends, suggesting entire chromosomes (Figure 1). The 18S ribosomal coding DNA sequence (rDNA ITS) has been identified on the scaffold ESM_6 between 2,312kb and 2,322kb, in a locally enriched transposable elements (TEs) region of 100kb (20 TE copies, see TE section below, Figure 1). This region was covered at 2,292X due to the tandem duplication of the element that has been assembled in a single copy. The mating-type locus (MAT1-2-1 allele, GenBank accession: AF380364.1, McGuire et al. 2001) was located at the end of the scaffold ESM_7 (Figure 1).

**Figure 1.**
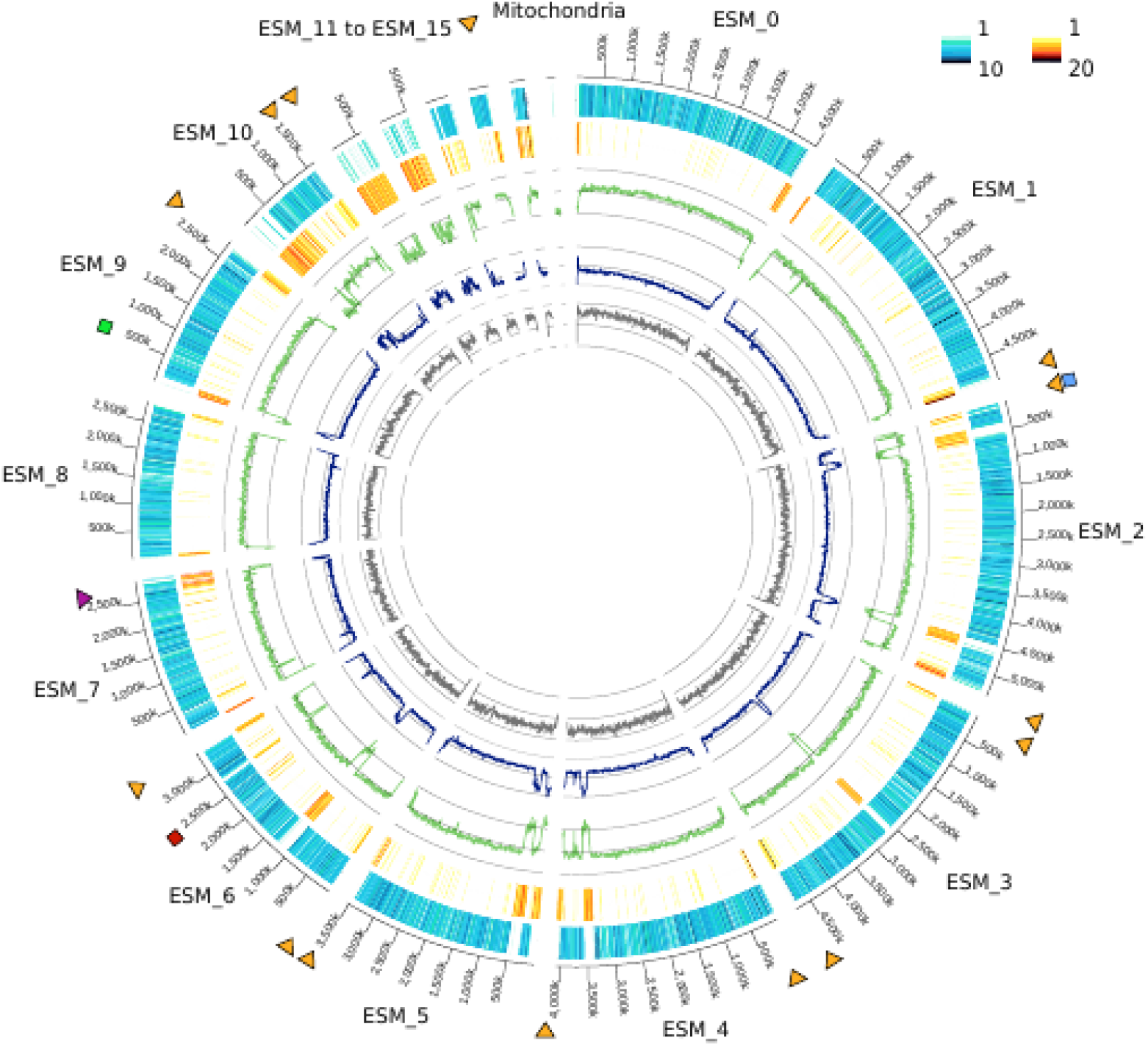
Circular representation of the final genome assembly of the *Cryphonectria parasitica* isolate ESM015 First circle: ID scaffolds, telomeric motifs (orange triangle), mating type locus (dark purple triangle), genes mapped in Eseubio-Cope et al. (2009) are nam-1 (blue square), rDNA ITS (red square) and β-tubulin (green square). Second circle: Number of predicted genes using BRAKER (Hoff et al. 2019) per 10kb. Color scale is shown in the top-right. Third circle: Number of transposables elements per 10kb. Color scale is shown in the top-right. Fourth circle: GC rate per 10kb, black lines represent 60%, 50%, and 40%. Fifth circle: Illumina reads mapping coverage. Sixth circle: Nanopore reads mapping coverage. Black lines represent 0, 30 and 60X. Coverage of the mitochondria and the 20kb ITS rDNA region on the ESM_6 scaffold has been excluded for clarity.

**Table 1 :** Assembly statistics for the genomes of the *Cryphonectria parasitica* isolates EP155, YVO003 and ESM015

| Reference | EP155<br>Crouch *et al.* 2020 | YVO003<br>Demené *et al.* 2019 | ESM015<br>This study |
| --- | --- | --- | --- |
| Assemblers | Arachne | PbcR | HybridSPAdes |
| Data type | Sanger | PacBio and<br>IonTorrent | ONT and<br>Illumina |
| Coverage (x) | 8.54X | >100X | 94.7X |
| GC-count (%) | 50.8 | 52.9 | 50.9 |
| Contigs | 26 | 35 | 18 |
| Assembly length (bp) | 43,870,411 | 39,261,148 | 43,305,322 |
| N50 (bp) | 5,118,729 | 2,722,809 | 4,077,748 |
| L50 (#) | 4 | 6 | 5 |
| N90 (bp) | 1,008,593 | 170,449 | 2,847,727 |
| L90 (#) | 11 | 26 | 10 |
| Max length (bp) | 7,437,273 | 4,766,472 | 5,381,768 |
| Min length (bp) | 2,579 | 19,618 | 39,291 |
| Complete BUSCO (%) | 98.6 | 98.7 | 98.7 |

### Compartmentalization and gene landscape of the two reference genomes

Gene prediction and annotation of the ESM015 genome assembly identified 12,015 gene models (Table S4). From these 12,015 models, 95% (n=11,461) were predicted to contain at least one protein domain on the different Interpro databases tested (Table S4), and 63% (n=7,564) show homology with a protein registered in the manually curated SwissProt database (Table S4). Gene ontology (GO) assignment based on these Interpro and SwissProt databases was successful for 67.3% of the total gene number (n=8,088). SignalP prediction resulted in 1,081 genes with a putative signal peptide, while EffectorP identified 1,117 putative effectors. The combined predictions both with a likelihood greater than 0.5 yielded 88 CSEP (Table S4). A total of 47 predicted secondary metabolite (SM) clusters, containing between 5 to 53 genes, were annotated using antiSMASH. Near half (n = 23) of these clusters were categorized in the Type 1 polyketide synthase group. Finally, 3.1% of total gene number (n=372) failed to be assigned to any known protein domain or to any homologous proteins of known function (Table S4). Genes were separated by an average intergenic distance of 1.9kb (SE = 0.034). Only few genes (n=318) were separated by large intergenic distances indicating a low genomic compartmentalization (Figure 2). The 88 putative CSEP had a similar intergenic distance than the other genes (1.87 kb, SE=0.15, t-test, p-value = 0.44). Identical results were observed for both SignalP and EffectorP predictions (Fig. S2A and S2B). The SM clusters were located on all scaffolds, except for four small ones (ESM_11, ESM_12, ESM_15 and ESM_16), and were not preferentially located in gene sparse regions (Fig. S2C).

**Figure 2.**
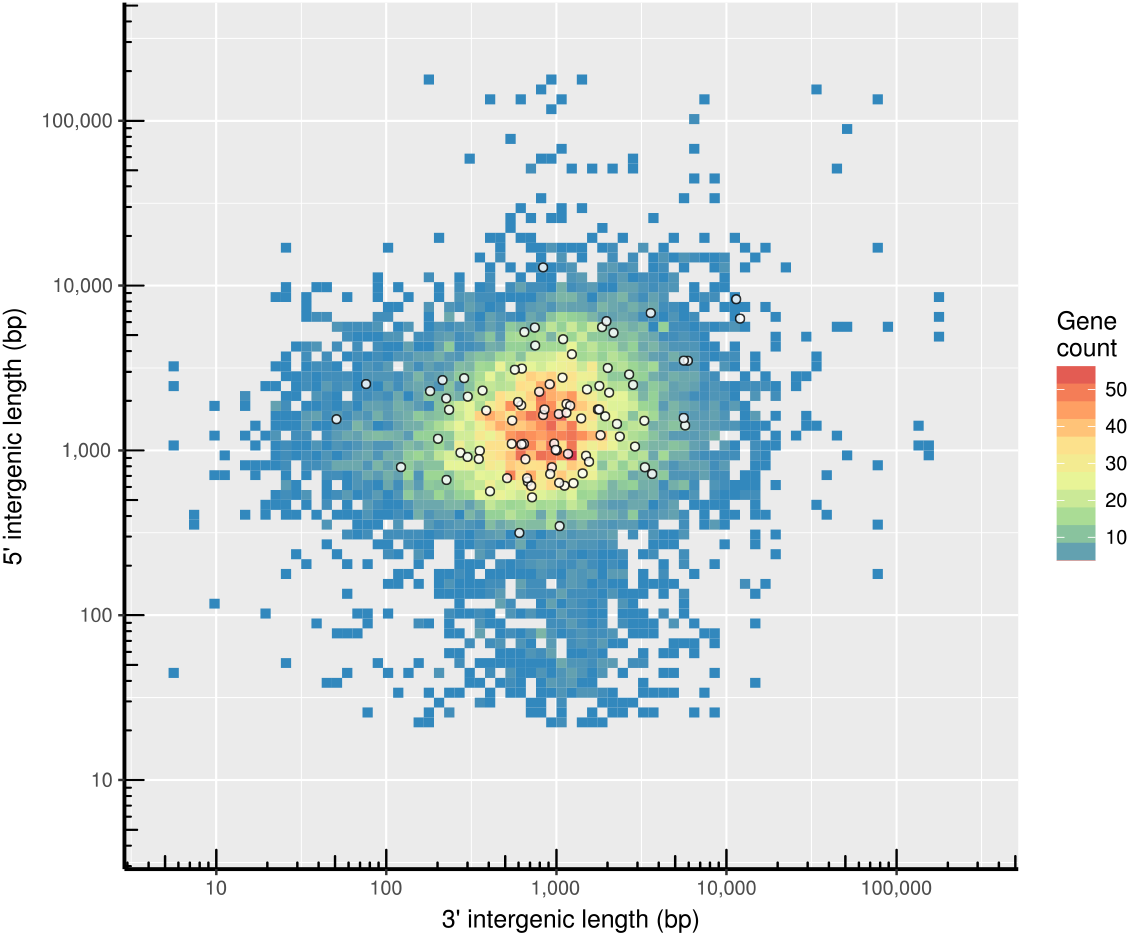
Estimated intergenic distances between the predicted genes of the ESM15 genome. Density of genes for each pair of distances (3’ and 5’ intergenic distances) is given according to the legend on the right. White circles are the candidate secreted effector-proteins predicted (CSEP) using the combined predictions with a likelihood greater than 0.5, of EffectorP (Sperschneider et al. 2018) and SignalP (Armenteros et al. 2019).

Out of the 12,015 predicted genes, 92% (11,056) had at least one orthologous predicted in the EP155 reference genome (Crouch et al. 2020; Table S5). After querying the missing gene models of ESM015 against the EP155 genome assembly with Blastn, 611 (5.1% of all genes) of them were found with 100% of identity on their full length, and 308 (2.6%) had minimum 80% of identity on at least 80% of the alignment length. Only 40 predicted genes of the ESM015 genome were considered lost in EP155, due to a high divergence (sequence similarity smaller than 80%), an alignment overlap < 80%, an absence of the start codon, or a complete absence of the coding sequence (i.e. no hit: 3 genes). Considering these 348 lost or partial genes, most of them (269; 77.3 %) had not known function (i.e. no annotated GO function; Table S5). Eleven of these lost or partial genes were found in different secondary metabolites clusters, and two were predicted as CSEP. Conversely, out of the 11,609 genes predicted in EP155 (Crouch et al. 2020), 94.2 % (10,932) had at least one orthologous, 1.1 % (133) were found in the ESM015 sequence but not predicted as a gene model, and 1.92 % (223) were divergent (between 80 and less than 100% of sequence similarity), or with an incomplete overlap (between 80 and less than 100%) (Table S6). Finally, 321 (2.7%) predicted genes in EP155 were considered lost in ESM15 generally because of a missing start codon. Only 11 were considered totally absent (no hit after searching in the ESM015 genome sequence with blastn). Out of the 321 genes considered lost in ESM015, 279 (86.9%) had no annotated GO function, and nine were found in a SM cluster in EP155. Most of these lost genes were located within or close to a TE cluster (Fig. S4).

### Comparison of the TE repertoire between the two genomes and identification of RIP signatures

The transposable element database, curated and representative of both ESM015 and EP155 genomes, contained 18 TEs consensus sequences (9 RNA TEs, 7 DNA TEs, one potential host gene and one unclear RNA/DNA TE). Annotation of the ESM015 and EP155 genomes led to 1,915 complete and incomplete TE copies (total length of 4.37Mb, 9.97% of the genome) and 2,116 copies (total length of 4.99 Mb, 11.37%; Table 2) respectively. Class I retrotransposons were the most abundant TEs, counting respectively for 9.48% and 10.79% of each genome, while class II transposons represented only 0.38% and 0.44% (Table 2). The ESM015 assembly contains an average of 4.6 TEs per 100kb (CI95%=0.82), which are not randomly distributed along the genome (number of TE per 100kb, χ2=259.46, df=5, p-value < 2.2e-16; Fig. S3). Those TEs were usually found clustered at the end and, sometimes, in central position of the scaffolds (Figure 1, Fig. S4).

**Table 2 :**
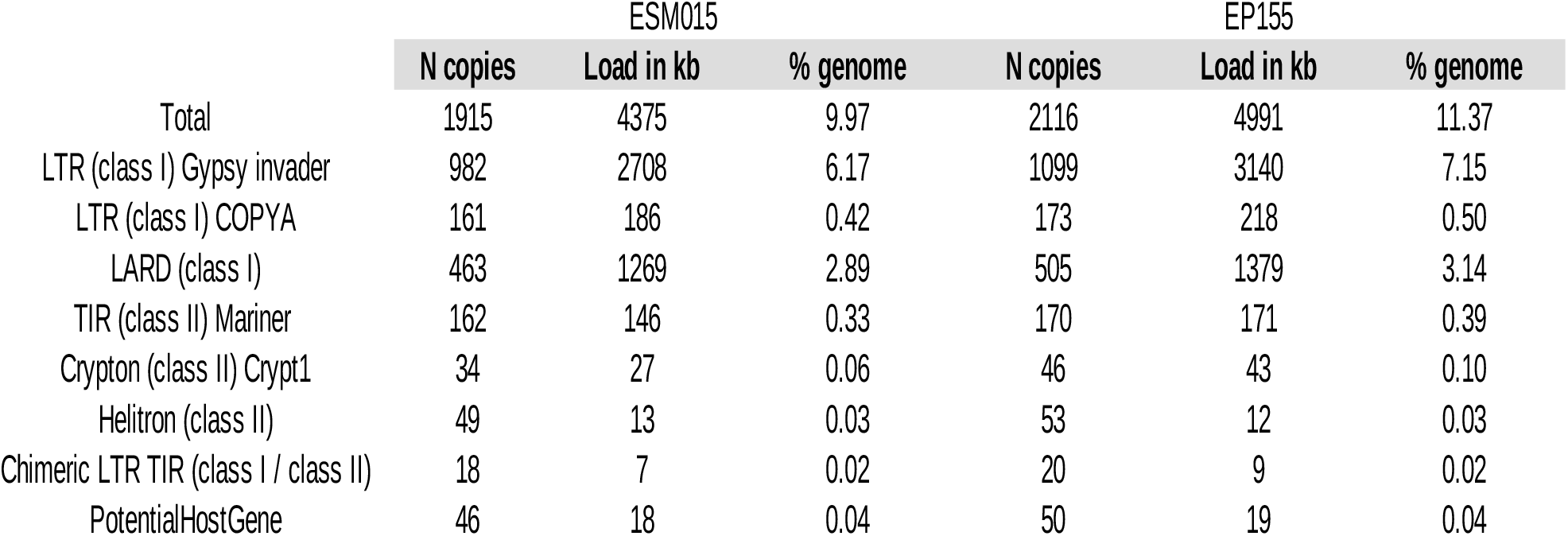
Distribution of the different categories of transposable elements in the ESM015 and EP155 genome assemblies

The most widely represented TE is a Gypsy like element (Class I), described as Metaviridae (Gypsy/Ty3 elements) in EP155 (Crouch et al. 2020), which contains the five protein coding domains and the long terminal repeats (LTR) sequences needed for its duplication (Wicker et al. 2007). Its cumulative genome coverage represented more than 2.7Mb (982 copies, 6.17% of the genome) in the ESM015 genome and 3.1Mb in the EP155 genome (1,099 copies, 7.15% of the genome, Table S7). The mating type locus, located on the scaffold ESM_7, is close to one of these regions rich in Gypsy elements (Figure 1), including fifty Gypsy copies out of the 95 incomplete or complete copies (270 kb total length). The difference in copies of TEs (mainly the Gypsy element) between the two genomes was likely due to the variability found in the mating-type region, sub-telomeric and telomeric regions that were all less well assembled (i.e., shorter) in the ESM015 than in the EP155 assembly. One identified Crypton DNA transposon was related to Crypt1, the only *C. parasitica* element that has been shown experimentally to be active (Linder-Basso et al. 2001). Five full length copies (FLC) of Crypt1 have been detected within the ESM015 genome and eight in the EP155 genome with a nucleotidic identity ranging from 95.5% to 99.8%.

Relationships between CSEP genes and TEs was investigated in ESM015 genome. The average minimal distance between TEs and genes predicted to be effectors by EffectorP (63.16kb, SE = 2.07kb) was slightly significantly different from the mean distance of all the other predicted genes (67.36 kb, SE = 0.76kb; Wilcoxon rank test, W = 5425810, p-value =0.034), but not for secreted proteins predicted by SignalP (66.27 kb, SE = 2.23kb; Wilcoxon rank test, W = 5115078, p-value = 0.49) (Fig. S5). By contrast, secondary metabolites (SM) clusters were found highly significantly closer to TEs than the other predicted *C. parasitica* genes (average minimal distance 46.03kb, SE = 1.96kb; Wilcoxon rank test, W = 3657390, p-value = 4.009e-15, Fig. S5). Out of the 47 SM clusters predicted, ten were located inside a TE-rich region. Most of the nucleotide sequence divergences between TE copies and their consensus sequences was estimated between 15 and 25% in the ESM015 genome (Figure 3A). We did not detect any significant differences between the proportions of TE families identified between the two genomes (Table 2, Table S6). The presence of several AT-rich isochores (i.e. with a GC content <50%; Fig. S6) was also identified in some regions of the ESM015 genome. The detection of RIP activity showed different results depending of the RIP index used. According to the pre-RIP index, no RIP signature could be revealed for both the two genomes and the four tested groups of TEs (Crypt1; all DNA TEs, the Gypsy family and the remaining RNA TEs. Nonetheless, estimates of the two “post-RIP” indexes for the four classes of TEs were either significantly higher or lower than the two estimated baseline frequencies (0.67 and 0.82 for TpA/ApT and CpG/GpC respectively, Figure 3B), suggesting a RIP activity for both the two genomes.

**Figure 3.**
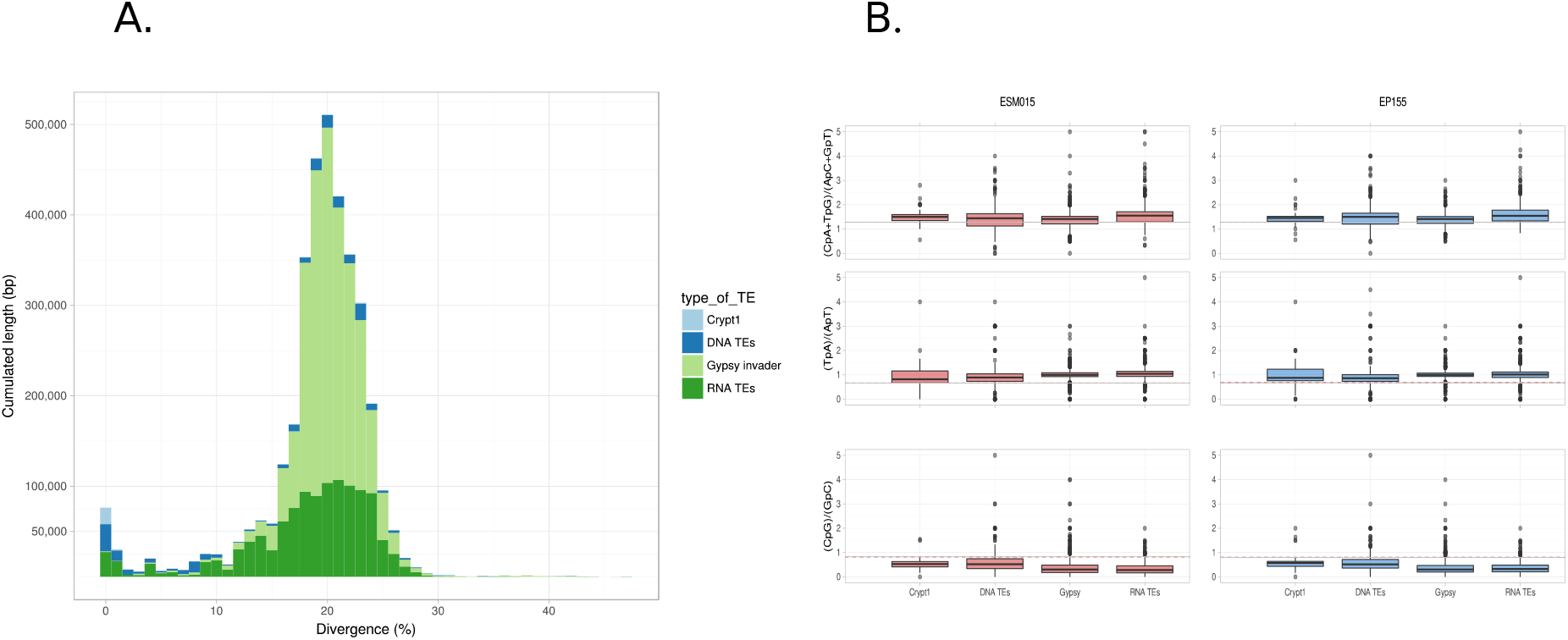
Transposable elements (TEs) landscape of the ESM015 and EP155 genome. **A.** Nucleotidic divergence of TEs copies to their consensus, grouped by type of TEs. **B.** Inference of the signature of the repeat induced point (RIP) mutation. Three estimated indices are presented (CpA+TpG)/(ApC+GpT), TpA/ApT and CpG/GpC (see text for details). The dashed red line indicates the threshold values obtained from the genomic sequences without TE for the corresponding genome (ESM015 or EP155). Confidence intervals at 99.9% for the threshold values are indicated in grey.

### Major structural variations between the two genome assemblies

Alignment length between the two genomes was 43.34Mb, with a mean nucleotide identity of 95.1%. This pairwise alignment revealed nine major synteny breaks (i.e. genomic translocation or inversion > 100 kb; Figure 4). Out of these nine synteny breaks, six were inter-scaffold translocations involving seven scaffolds of EP155 (scaffolds 1, 2, 5, 6, 7, 10 and 11) and five for ESM015 (ESM_2, 4, 5, 7, 10) (Figure 4, details in Table S8). The three remaining major intra-scaffold translocations involved one extremity of the scaffold 2 of EP155 and ESM_11 and ESM_12 of ESM015, and the scaffold 6 of EP155 versus the scaffold ESM_6 of ESM015. The two detected major translocations on the scaffold 2 of EP155 are located in a region rich in TEs of about 1.7Mb between the MAT locus and a telomeric region (Fig. S7).

**Figure 4.**
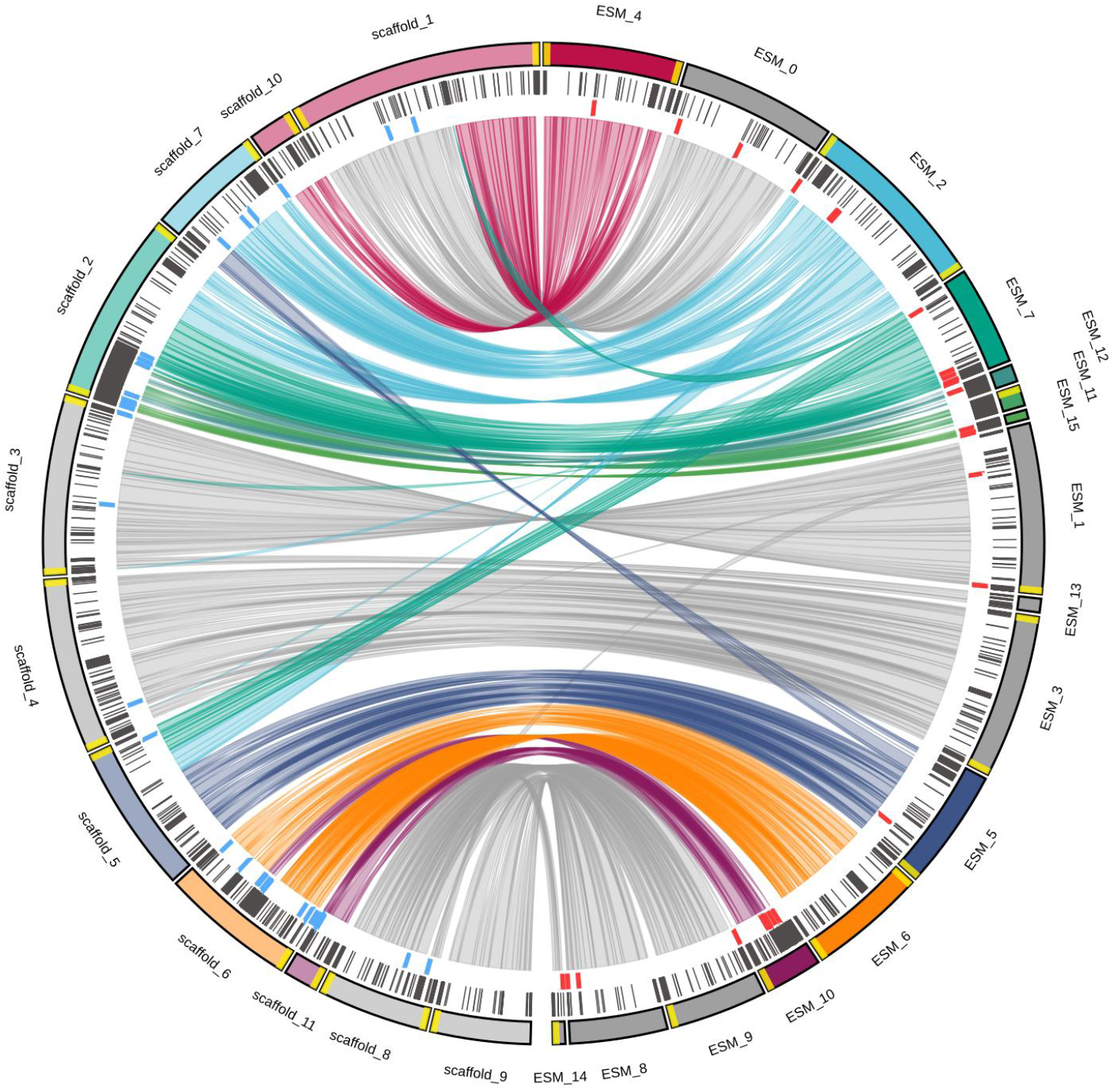
Syntenic relationships and genomic rearrangements between the two *Cryphonectria parasitica* genomes of the isolates ESM015 and EP155. Circular representation of the syntenic relationships between the two genomes. Links are the syntenic regions higher than 10kb and with a similarity higher than (95%). Coloured scaffolds are scaffolds presenting chromosomal rearrangements between the two genomes. Yellow marks at the extremities of scaffolds are the identified telomeric motifs. Black marks of the second circle represents the density of TEs. Blue and red marks represent the location of the Crypt1 copies for EP155 and ESM015 genomes respectively.

All the detected translocations were associated with one break per scaffold, except for the scaffolds 2 and 5 of EP155, and the scaffold ESM_2 of ESM015 with two breaks. All these rearrangements were confirmed by the visual inspection of reads mapped on the two assemblies. In the regions of the detected synteny breaks, Nanopore long reads generated in this study fully mapped on the new de novo ESM015 genome assembly. However, in these same regions, they only mapped for one side on the previous published EP155 genome assembly (Fig. S8). Conversely, the Sanger reads generated in Crouch et al. (2020) fully mapped on the EP155 assembly for these regions, but not on the ESM015 genome assembly (Fig. S8). These results suggested that the identified chromosomal rearrangements were not chimeric assemblies for the two genomes. At the exception of the rearrangement between the scaffold 6 of EP155 and ESM_10 of ESM15, and associated with the two class of TEs, Crypt1 and Crypt2 (Figure 4), chromosomal translocations were linked to regions poor in TEs and rich in predicted genes (Figure 4 and 5A). We observed that breaks are not included within homologous regions between the involved scaffolds (Figure 5A).

**Figure 5.**
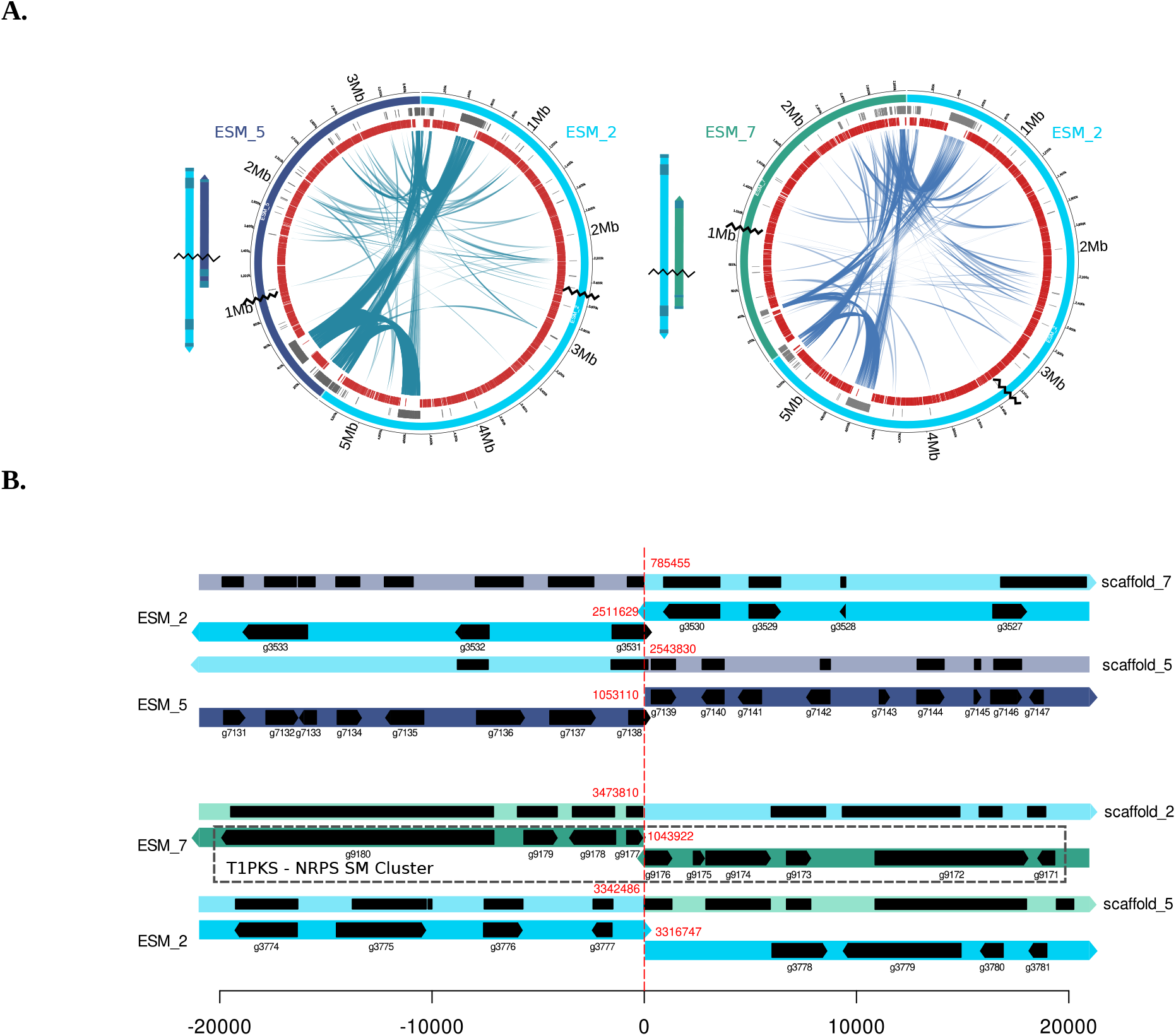
Zoom on the syntenic relationships between the three scaffolds of ESM015 involved in two major inter-chromosomal translocations. **A.** Circular representation of the ESM015 scaffolds involved in the inter-chromosomal translocations. Transposable elements are shown in grey and genes in red. Synteny breaks are represented by the black broken lines. All syntenic links obtained with Nucmer are shown. **B.** Schematic representation of the 40kb genomic regions around the chromosomal breaks for the three translocation events. Positions of the break for each scaffold are given in red, and ID of genes and their position on the scaffolds are in black. The secondary metabolite cluster T1PKS is indicated by the rectangle in dotted line on the scaffold ESM_7 of the ESM015 isolate.

These results suggested that these chromosomal rearrangements were associated with non-homologous end-joining events. Focusing on the four synteny breaks involving ESM_2, ESM_5 and ESM_7, three events would have led to gene disruptions, for which functional prediction were a transcription factor (g3531.t1 on scaffold ESM_2), and two MFS transporter (g7138.t1 scaffold ESM_5 and g9176.t1 on scaffold ESM_7; Figure 5B). Finally, the synteny break between the ESM_7 of ESM015 genome and the scaffolds 5 and 2 of the EP155 genome occurred in a predicted SM cluster (T1PKS) (Figure 5B), a class of polyketide synthase putatively involved in mycotoxin and phytotoxin productions (Bills and Gloer 2016).

## Discussion

### Long reads Nanopore sequencing technology: an efficient method to obtain a near-chromosome level genome assembly in fungal species

This study confirmed that long reads sequencing produced with Nanopore technology is well adapted to assembly small genomes such as found in fungal species, and for regions with low complexity (Rajarammohan et al. 2019). Our final assembly was close to a near-chromosome level assembly. Its total length was 43.3Mb, close to the 43.8Mb of the EP155 genome assembly (Crouch et al. 2020), and contained four putative chromosomes with both telomeric regions and six scaffolds with one telomeric region. This result is consistent with the number of chromosome and genome size estimated in a previous cytological and electrophoretic karyotyping study (i.e. 9 chromosomes and 50 Mb estimated in Eusebio-Cope et al. 2009). The large quantity (66.0 μg) DNA obtained using the new DNA extraction protocol for fungal species was a key of this success. This protocol resulted in a DNA extraction highly concentrated in double-stranded DNA, which allowed to perform several steps of purifications and size selection. Another significant advantage of this protocol is the avoidance of using heavy and time-costly methods such as centrifugation on a caesium gradient and numerous purification steps (Cheeseman et al. 2014, Demené et al. 2019). The use of this new DNA extraction method should greatly help the assembly of new genomes at the chromosome level in other filamentous fungi.

In addition, the comparison of few methods of assembly was very useful to assess a valuable strategy to obtain a near-chromosome assembly without generating chimeric scaffolds. The comparison of three methods, and the estimate of various metrics, beyond the classic L50 and N50, greatly helped to identify the less fragmented and chimeric assemblies. HybridSPAdes (Antipov et al. 2016) is a hybrid method based on de Bruijn graph algorithm (the two others used in this study were based on overlap-layout-consensus, OLC, method). Associated with short reads and the npScarf scaffolding tool, it was considered as the most satisfactory method for assembling this genome using the recent Nanopore technology sequencing. In addition, our study confirmed the importance of performing several steps of manual curation, and identifying possible chimeric scaffolds (Courtine et al. 2020). However, several repeated regions along the genome could still be imperfectly assembled (estimated to 0.5 Mb, a little more than 1% of the genome). This result called for additional studies to better identify advantages and drawbacks of each method, according to the size, the genome content, and the sequencing technology used (Lu et al. 2016, Bayat et al. 2020).

### Stable transposable elements and CSEP repertoires between the native and the introduced *C. parasitica* genomes

The proportion of TEs in the ESM15 genome was estimated to 10%, similar to the estimates performed on the EP155 reference genome (Crouch et al. 2020). More than two-third of TEs belong to the Gypsy family, the most abundant TE in the fungal kingdom (Muszewska et al. 2011). A high genetic divergence among most TEs sequences was estimated, suggesting ancient genomic duplications and subsequent accumulation of mutations. This high divergence could be due to an ancient burst of the main TE families which was stopped by a molecular mechanism such as RIP mutations, described in fungal species (Selker et al. 1987). Contrary to Crouch et al. (2020), we identified a signature of RIP mutation in the two *C. parasitica* genomes by comparing different dinucleotide ratio between TEs and other gene sequences. It was identified by estimating a higher TpA/ApT ratio in TEs, which is one of the products of RIP mutation (Margolin et al. 1998). By contrast, the more classical target considered in RIP analysis (CpA + TpG / ApC + GpT; Selker 1990) produced no robust proof of RIP signature for most TE families, as reported in Crouch et al (2020). However, the targets of RIP may differ significantly between fungal species (Ikeda et al. 2002). For example, the analysis of the Gypsy elements showed that CpG was the preferred target of the RIP mutation in *C. parasitica* (Clutterbuck 2011). By estimating a new index (CpG/GpC), depletion of this RIP target was identified in the two reference genomes for all TE families. Furthermore, a homologous protein of the RID gene of *Neurosporora crassa* (GenBank: AAM27408.1), the only gene experimentally proven to be necessary for RIP activity (Freitag et al. 2002), was identified in *C. parasitica* genome (blastp e-value 8e-96, g9609.t1 with the same PFAM motif PF00145 known to be involved in RIP activity; Amselem et al. 2015). This supports that the RIP is active and repressed the TEs expansion in the genomes of *C. parasitica*. In addition, regular sexual reproduction that is necessary for the occurrence of RIP mutations (Selker 1990), has been described, or inferred, several times, although at different frequencies, in native and introduced areas in *C. parasitica* (e.g., Milgroom et al. 1996, Dutech et al. 2012, Demené et al. 2019).

We also detected a slight gene compartmentalization in the *C. parasitica* genome, and, at the exception of secondary metabolites, no clear association between genes and TEs. Transposable elements were often found clustered in telomeric and sub-telomeric regions in *C. parasitica*, as well as the secondary metabolite (SM) clusters. The coding sequences predicted to produce effectors were slightly located closer to TEs than other genes, and no particular proximity with TEs has been found for putative secreted protein coding sequences. The concept of “two-speed genome”, assuming that genes involved in host-pathogen interaction like CSEPS, would be clustered together in regions having a rapid evolution due to the presence of active TEs in their neighborhood, is indeed widely debated in fungi (Frantzeskakis et al. 2019, Torres et al. 2020). Our analysis, more consistent with a low gene compartmentalization and stable CSEP and SM repertoires, is in agreement with Stauber et al. (2021) who have sequenced a hundred of *C. parasitica* samples in Europe using short-reads Illumina. These results have also been described in other plant fungal pathogens of natural ecosystems (Hartmann et al. 2018), and also for other fungal species infecting crops (Stam et al. 2018, Schwessinger et al. 2018).

Comparison of the gene landscape between the two genomes also showed no major variation. Most of the predicted genes in one genome were present in the other one. The small number of missing genes either had unknown functions or did not have a start codon which may be due to sequencing or annotation errors. More noteworthy, we did not detect any significant changes in the SM and CSEP gene repertoires between the two genomes. This small variation in gain and loss of genes confirmed the results obtained in Stauber et al. (2020, 2021) on the hundred of *C. parasitica* isolates sequenced with Illumina. Similar results were also obtained in *Microbotryum sp*., the causal agents of anther-smut diseases, for which the low number of gene gains and losses observed between a ten of pairs of isolates have been considered neutral or lowly deleterious (Hartmann et al. 2018). Large variations in gene repertoires, especially associated with effector gene candidates have been regularly described within fungal pathogen species or within genera infecting different host species, and considered as a signature of host specialization (e.g., Yoshida et al. 2016, Wang et al. 2020). Such a signature was not detected between the EP155 genome infecting the American chestnut (*C. dentata*), and ESM015 infecting Asian chestnuts (such as *C. mollisima* and *C. crenata*). However, molecular interactions between *C. parasitica* and the Chestnut species may be more quantitative, and/or plastic, than qualitative, involving other metabolic pathways and regulations (Lovat and Donnelly 2019). Furthermore, the non-significant variation in host-virulence observed between native and introduced strains of *C. parasitica* against the European Chestnut also suggested that no or low selective pressure has affected the virulence traits of the fungus during the European introduction (Dennert et al. 2019).

### Numerous inter-chromosomal translocations with putative important consequences on sexual reproduction of *C. parasitica*

We detected six major inter-chromosomal translocations between the two genome assemblies, involving four scaffolds, and in some cases, comprising two events on the same chromosome. We cannot totally rule out that chimeric assemblies may explain the detected chromosomal rearrangement between the two de novo genome assemblies. However, we first observed that the genome assembly produced in this study for the ESM015 isolate had no identified major synteny breaks with the assembly produced in Demené et al (2019), using another isolate (i.e., YVO003, an isolate sampled in France), and another sequencing technology (i.e. Pacific Bioscience technology). This congruence between the two de novo genome assemblies, added to the fully mapping of Nanopore long reads on the new genome assembly in the assumed regions of synteny breaks with the EP155 genome, gave a high credit on our results. Similarly, we checked the mapping of Sangers reads produced in Crouch et al. (2020), and observed that several of these reads passed through the regions of synteny breaks for the EP155 genome assembly (Fig. S8). In addition, this EP155 assembly was performed using several plasmid/fosmid subclone sequencing, targeted sequencing of genomic gaps and was validated by a genetic linkage map (Crouch et al. 2020). These results should discard the possibility of several chimeric assemblies, especially in the regions with no repeated element. By contrast, a small number of large intra-chromosomal inversions and duplications were detected from this genomic comparison, except close to the mating-type region. These differences in extend between the intra- and inter-chromosomal variations were less often described in the genomic studies at the species level; chromosomal inversions, duplication or a limited translocation of genes being more abundantly described in fungi (e.g. Chuma et al. 2011, Hane et al. 2011, Raffaele and Kamoun 2012, Plissonneau et al. 2018). Inter-chromosomal translocation were however several times identified between species of the same genus (e.g. Wang et al. 2020), between isolates of mainly asexual species (de Jonge et al. 2013, Olarte et al. 2019, Tsushima et al. 2019), but also in few cases, in partially sexual species, such as *Saccharomyces cerevisiae* (Hou et al. 2014, Chang et al. 2013). This low number of reports on inter-chromosomal rearrangements may be explained by still a limited number of comparisons of genome assemblies at the near-chromosome level. We assume that the new sequencing technologies, such as those used in this study, will allow to obtain rapid and robust assemblies at the chromosome level for numerous genotypes within species, and would lead to more frequent report of such events in fungal species in a next future.

The synteny breaks were not associated with homologous regions between the involved scaffolds, especially with no cluster of repeated sequences, such as TEs, in the proximity of the breakpoints. These results suggest that these inter-chromosomal translocations were associated with random double-break strands (DBS) of the chromosomes, and led to the repair of these DNA breaks by non-homologous end-joining events. This process has been described in several Eukaryotes, and especially in fungi (e.g. Hou et al. 2014, Yadav et al. 2020). Double break strands may occur during episodic stresses, for example during some specific recombination events (Yadav et al. 2020), and have been described under laboratory conditions (e.g. Dunham et al. 2002), and also under natural conditions (e.g. Chang et al. 2013). For *C. parasitica*, we cannot identify which conditions may have led to this extensive chromosome shuffling between the native ESM015 and the introduced EP155 genome. Additional comparisons of *C. parasitica* genomes sampled in the native and the introduced areas must be performed to reconstruct the origin of these chromosomal rearrangements. Our two genome assemblies (Demené et al. 2019, this study), suggested that the North-American EP155 strain would not be the standard karyotype of *C. parasitica*, as argued in Eusebio-Cope et al. (2009). Furthermore, the identification of another karyotype in this later study (i.e. the GH2 strain sampled in Michigan) also suggested that chromosomal rearrangements would have occurred several times during the world *C. parasitica* invasion.

One of the consequences of these chromosomal rearrangements may be dramatic phenotypic changes among genotypes (e.g. Chang et al. 2013, Olarte et al. 2019). As discussed above, chromosomal rearrangements did not lead to major changes in the repertoire of genes between the two genomes. Nevertheless, the detected synteny breaks showed the disruption of three genes (one being included in a secondary metabolite cluster), and for which the phenotypic expression would be interesting to investigate. Chromosomal translocation may also change gene expression via, for example, change in DNA methylation or in proximity with some TEs (Castanera et al. 2016). The recent identification of changes in epigenetic marks between *C. parasitica* strains along the European gradient of colonization (Vukovic et al. 2019) provide more rationale for investigating the role of these mechanisms in the putative adaptation of the species to these new environments. Another important consequence of these chromosomal rearrangements is the impact on the success of sexual reproduction between these genomes, and the raise of putative pre-zygotic barriers, leading to incipient speciation (Hou et al. 2014, Yadav et al. 2020). Inter-chromosomal translocations lead to abnormal segregation of homologous chromosomes during meiosis (Kistler and Miao 1992), and possible acquisition of an incomplete repertoire of genes in the progenies of parents with different karyotypes (Yadav et al. 2020). Pre-zygotic barriers involving these rearrangements may actually explain the limited number of intermediate genotypes among the predominant genetic lineages observed in south-western Europe, and despite the regular presence of the two mating types within populations (e.g. Dutech et al. 2010). Usually, chromosomal rearrangements are strongly reduced in sexual species (Plissonneau et al. 2018). However, the specific context of introductions, associated with genetic bottleneck and new environmental conditions, and the occurrence of clonal reproduction in *C. parasitica* in introduced areas (Milgroom et al. 1992, Dutech et al. 2010) may have favoured the fixation of new genetic lineages in this invasive fungal species. With the emergence of long reads sequencing technology, this hypothesis can be now more easily investigated in the *C. parasitica* populations, and allows us to understand the role of large structural genomic variations in the adaptation of this invasive species.

## Data accessibility

All bash, python and R scripts used for performing these analyses, TEs databases, genes annotations, and supplementary figures are available on the “Portail Data INRAE: Chromosomal rearrangements but no change of genes and transposable elements repertoires in an invasive forest-pathogenic fungus” at https://doi.org/10.15454/UTIB8U. This Whole Genome Shotgun project has been deposited at DDBJ/ENA/GenBank under the accession JAGDFO000000000. The version described in this paper is version JAGDFO010000000. Raw sequence data and the ESM015 genome assembly are available on the bioproject PRJNA700518.

## Supplementary material

Supplementary material is available at https://doi.org/10.15454/UTIB8U

**Table S1**: Assembly statistics for the genome of the Cryphonectria parasitica isolate ESM015 after using different assemblers

**Table S2**: Assembly statistics of the ESM015 genome using HybriSPAdes according to the chosen kmer size

**Table S3**: Gene prediction and annotation of the mitochondrial contig of the Cryphonectria parasitica isolate ESM015 using MITOS2 (Donath et al. 2019)

**Table S4**: list of genes predicted in the ESM015 genome assembly

**Table S5**: Comparison of the predicted genes of ESM015 versus those predicted in the EP155 genome

**Table S6**: Comparison of the predicted genes of EP155 versus those predicted in the ESM015 genome

**Table S7**: Details of the annotated TEs in the two genome assemblies of the Cryphonectria parasitica isolates ESM015 and EP155

**Table S8**: Genetic similarity among scaffolds and detection of major (i.e., > 100kb) synteny breaks between the ESM015 and EP155 genomes

**Figure S1**: KAT plots of the ESM015 genome assembly

**Figure S2**: Estimated intergenic distances between predicted genes of the ESM015 genome

**Figure S3**: Distances (in nucleotides) separating the pairs of TEs detected in the ESM015 genome assembly

**Figure S4**: Circular representation of the missing genes after comparison of gene predictions in the ESM015 and the EP155 genome assemblies

**Figure S5**: Violin plots of distances between TEs and all pairs of predicted genes or pairs of different classes of predicted genes (effectors predicted by EffectorP or secreted proteins predicted by SignalP), and secondary metabolites (SM) predicted by AntiSMASH).

**Figure S6**: estimated GC content in the ESM015 genome assembly

**Figure S7**: Circular representation of the genomic region around the mating type and their syntenic relationships between the ESM015 and EP155 genome assemblies

**Figure S8**: Visualization of the mapping of the Nanopore (ESM015) or Sangers (EP155) reads on the ESM015 and EP155 genome assemblies around the major detected synteny breaks

**Supplementary Material 1**: Protocol of DNA extraction for Nanopore long-reads genome sequencing

**Supplementary Material 2**: Curation of the TE databases

**Supplementary Material 3**: Statistics of the DNA extraction of the Cryphonectria parasitica ESM015 isolate

## Acknowledgements

We thank Michael Milgroom for the yield of the Japanese ESM015 isolates; N. Feau, B. Llorente, M. Foulongne-Oriol and C. Robin for their precious comments and corrections on previous versions of the manuscript; J-F Flot, B Schwessinger and one another anonymous reviewer for their evaluation for PCI Genomics, B. Penaud, L. Duvaux and I. Lesur Kupin for their help on genome assemblies; the URGI team (Unité de Recherche Génomique Info) of Versailles (France) for their support and training sessions for the transposable element predictions and annotations. We thank L. Frantzeskakis for sharing his R script used and modified for generating some figures. MinIon sequencing was performed at the Genome Transcriptome Facility of Bordeaux (grants from the Conseil Regional d’Aquitaine n°20030304002FA and 20040305003FA, the European Union, FEDER n°2003227 and Investissements d’avenir, N°ANR-10-EQPX-16-01). This work was supported by the ANR-12-ADAP-0009 (Gandalf project). A. Demené was supported by a PhD fellowship from the Ministère français de l’enseignement supérieur - Université de Bordeaux.

Version 6 of this preprint has been peer-reviewed and recommended by Peer Community In Genomics (https://doi.org/10.24072/pci.genomics.100013).

## Conflict of interest disclosure

The authors of this preprint declare that they have no financial conflict of interest with the content of this article.

## Author contributions

AD extracted DNA, sequenced and assembled the genome, performed the data analyses and interpretation and wrote the manuscript. BL performed the gene content analysis and helped in data interpretation and writing the manuscript. CD conceived and supervised the study and wrote the manuscript. SC-A conceived the protocol of DNA extraction for long reads, and CB. did DNA purification after extraction and sequenced the genome.

## Notes

https://github.com/Ademene/CParasiticaGenome_scripts_and_data

### Competing Interest Statement

The authors have declared no competing interest.

### Summary of Updates

In PCI Genomics format

