## Supplementary material for "Chromosomal rearrangements with stable repertoires of genes and transposable elements in an invasive forest-pathogenic fungus": Supplemantary Figures: FigureS1_Demene2021_ReSub.pdf

**Figure S1**: KAT plots of the ESM015 genome assembly
a) HybridSPAdes assembly using kmer size of 127, b) HybridSPAdes assembly using kmer size of 127 after the scaffolding step, c) HybridSPAdes assembly using kmer size of 127 after the scaffolding step and the first manual curation, d) HybridSPAdes assembly using kmer size of 127 after the scaffolding step and the second manual curation, e) The final ESM015 assembly, f) The Miniasm assembly, g) the Miniasm assembly with a previous correction of long-reads by short-reads and h) the Ra assembly.

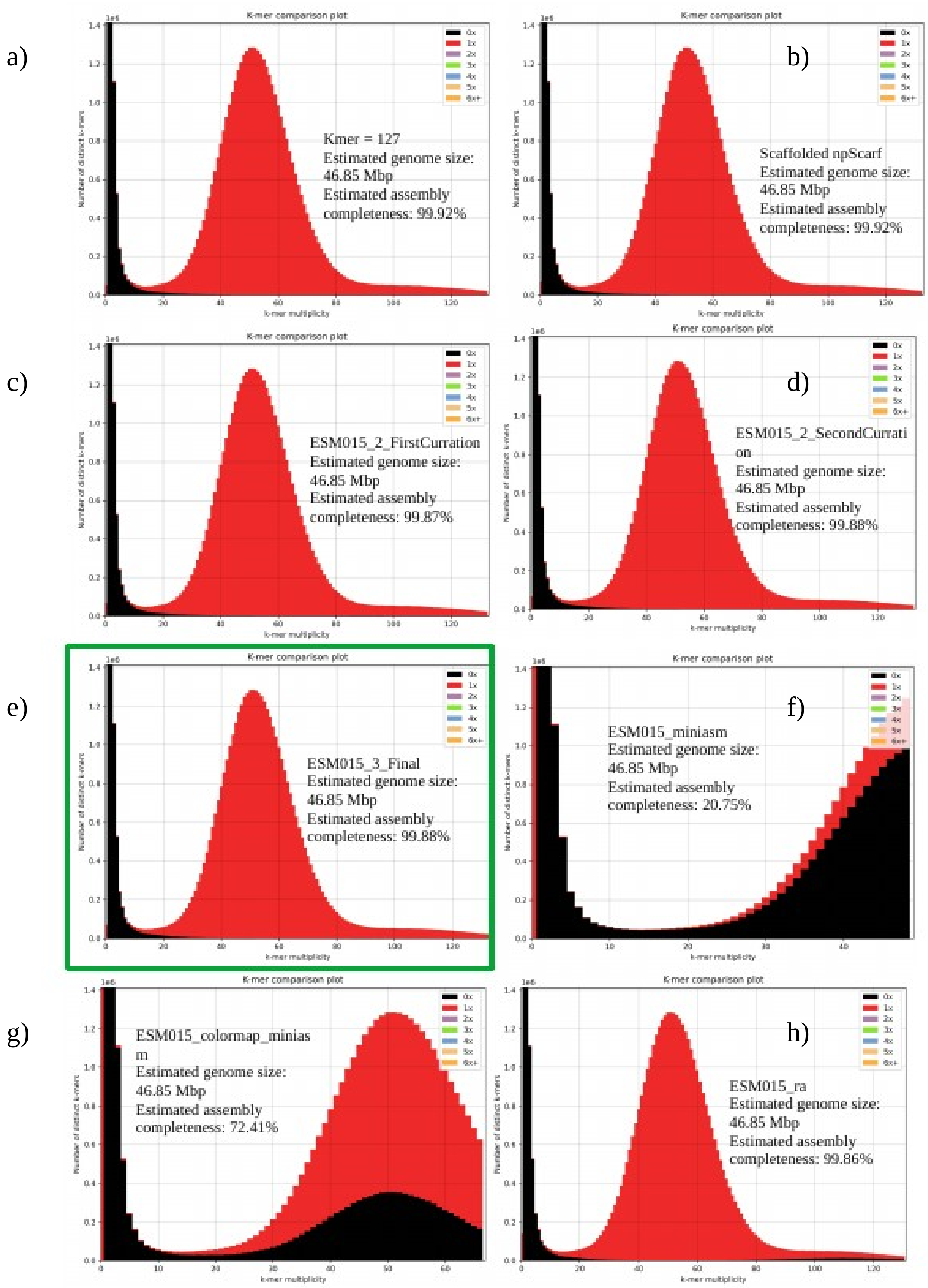
