## Supplementary material for "Chromosomal rearrangements with stable repertoires of genes and transposable elements in an invasive forest-pathogenic fungus": Supplemantary Figures: FigureS2_Demene2021_ReSub.pdf

**Figure S2 :** Estimated intergenic distances between predicted genes of the ESM015 genome
A. Yellow circles represent candidate secreted genes predicted by SignalP. B. Blue circles represent candidate effector genes predicted by EffectorP. C. Red circles represent candidate genes present in secondary metabolite clusters predicted by AntiSmash.

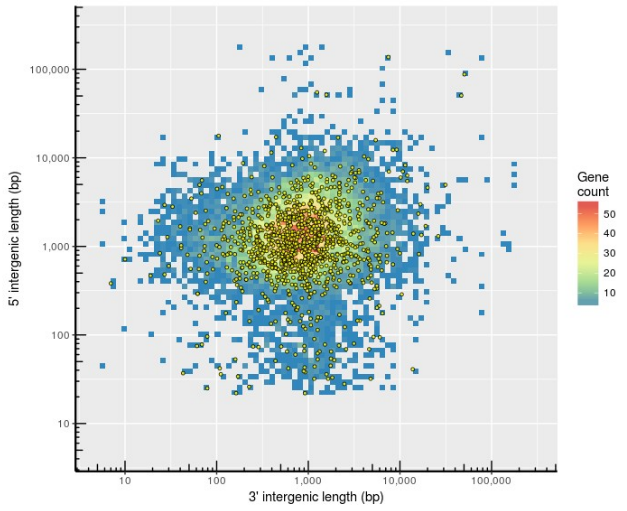


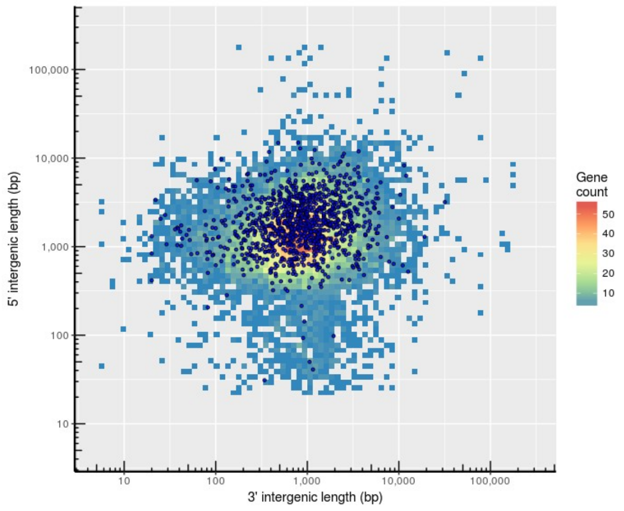


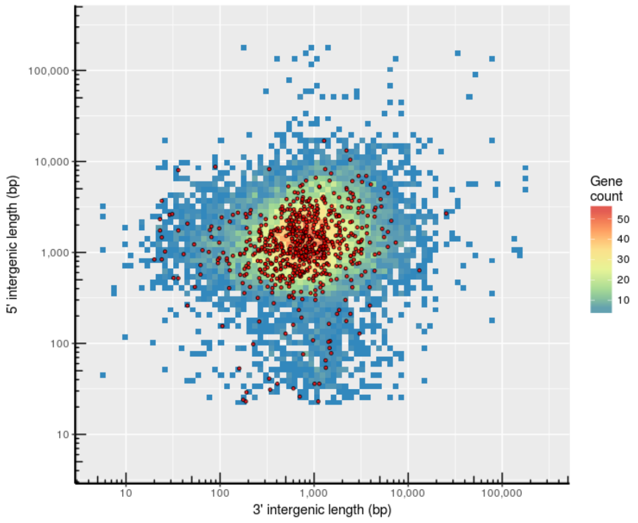
