## Supplementary material for "Chromosomal rearrangements with stable repertoires of genes and transposable elements in an invasive forest-pathogenic fungus": Supplemantary Figures: FigureS3_Demene2021_ReSub.pdf

**Figure S3** : Distances (in nucleotides) separating the pairs of TEs detected in the ESM015 genome assembly

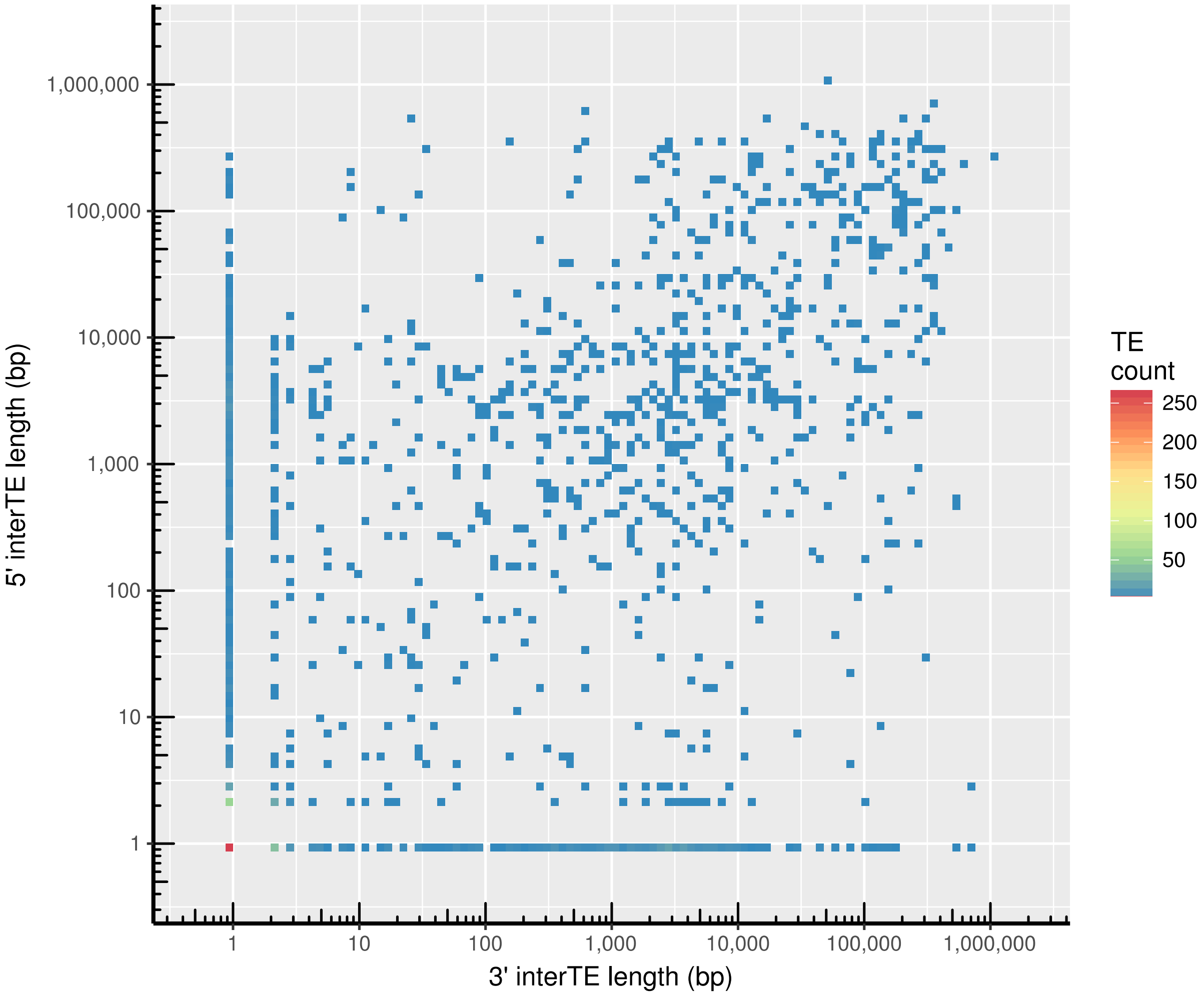
