## Supplementary material for "Chromosomal rearrangements with stable repertoires of genes and transposable elements in an invasive forest-pathogenic fungus": Supplemantary Figures: FigureS4_Demene2021_ReSub.pdf

**Figure S4:** Circular representation of the missing genes after comparison of gene predictions in the ESM015 and the EP155 genome assemblies

The ESM015 assembly is represented at the right, and EP155 at the left. Yellow rectangles represent the telomeric regions. The second circle in grey represents the detected TEs, on the third circle in red, the genes of the ESM015 genome missing in the EP155 genome, and in blue, the genes of the EP155 genome missing in the ESM015 genome.

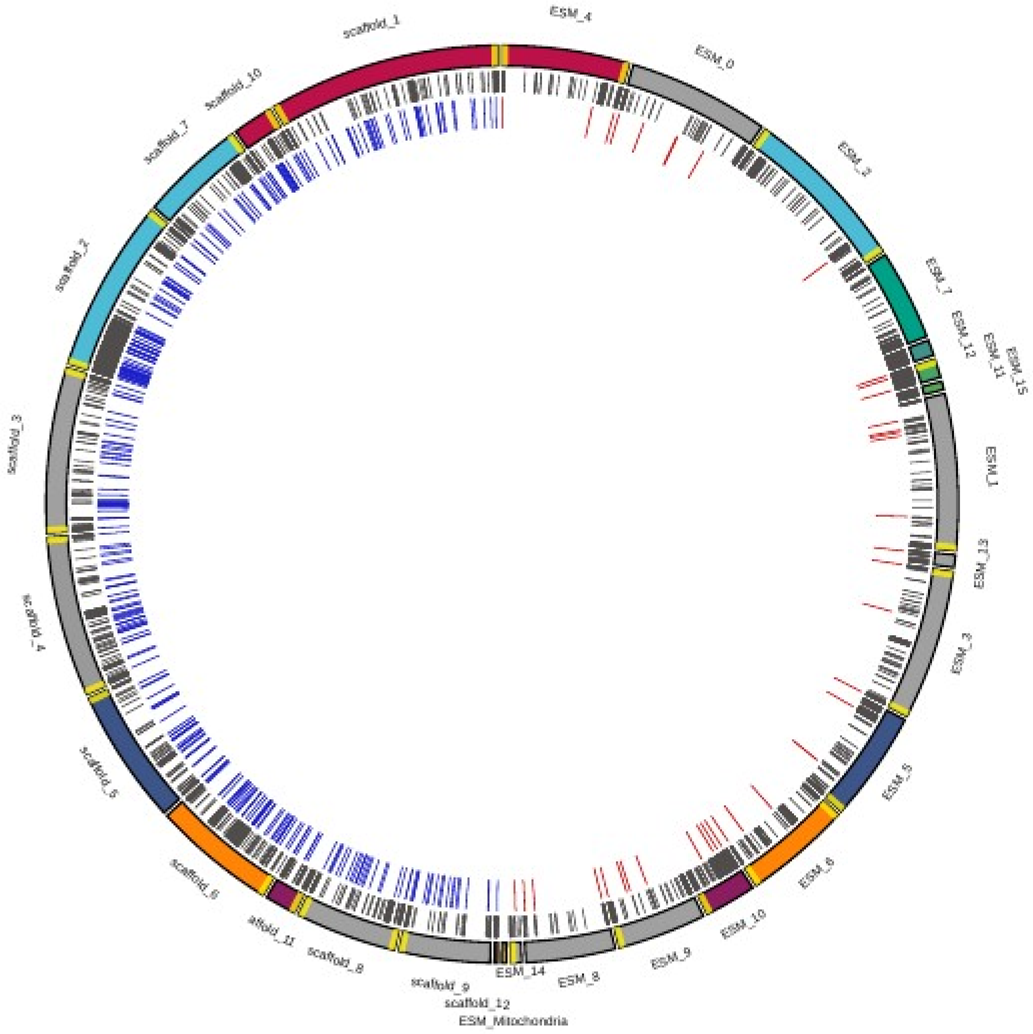
