## Supplementary material for "Chromosomal rearrangements with stable repertoires of genes and transposable elements in an invasive forest-pathogenic fungus": Supplemantary Figures: FigureS5_Demene2021_ReSub.pdf

Figure S5 : Violin plots of distances between TEs and all pairs of predicted genes or pairs of different classes of predicted genes (effectors predicted by EffectorP or secreted proteins predicted by SignalP), and secondary metabolites (SM) predicted by AntiSMASH).
Black points represent the mean distance of each class. Significativity of the Wilcoxon rank tests are indicated by NS (Not significant, p-value > 0.05), * (slightly significant, 0.05 > p-value > 0.01) or *** (highly significant, p-value <<< 0.001).

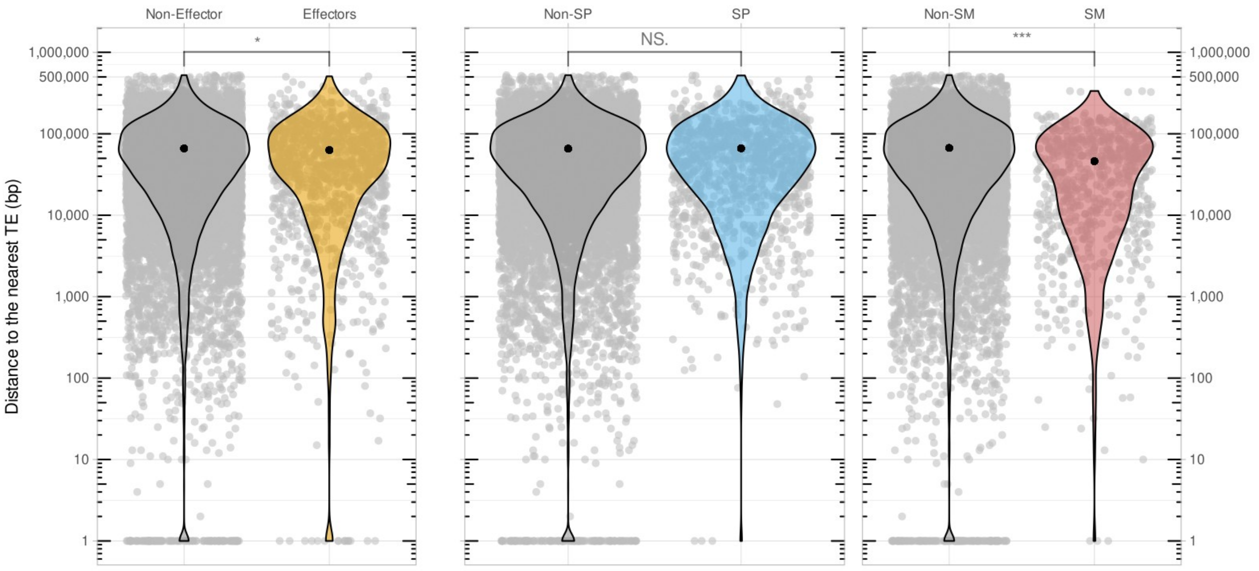
