## Supplementary material for "Chromosomal rearrangements with stable repertoires of genes and transposable elements in an invasive forest-pathogenic fungus": Supplemantary Figures: FigureS7_Demene2021_ReSub.pdf

**Figure S7 :** Circular representation of the genomic region around the mating type and their syntenic relationships between the ESM015 and EP155 genome assemblies
The EP155 genome assembly is represented at the left and ESM015 at the right. The mating type is represented by the red line.

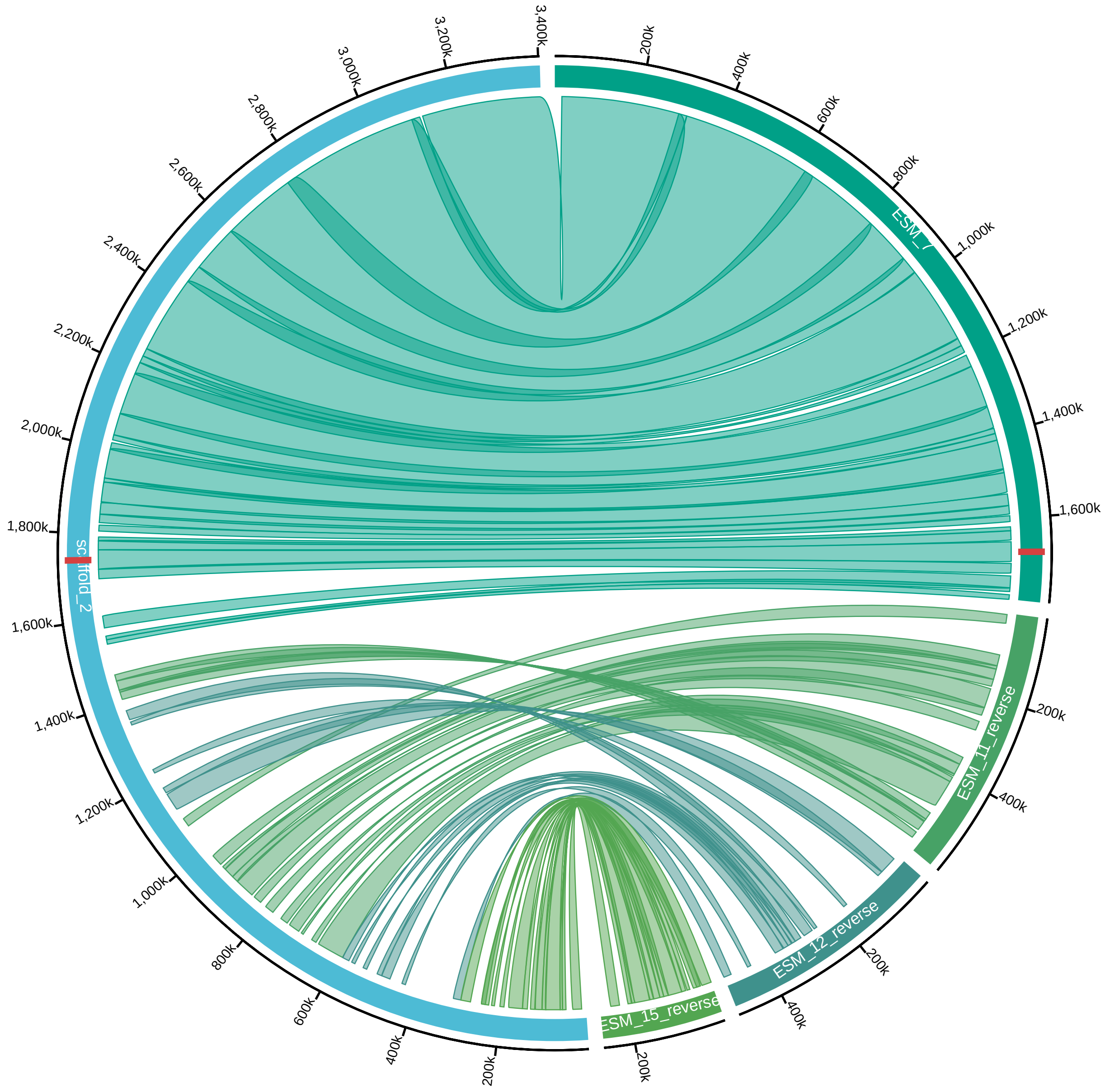
