## Supplementary material for "Chromosomal rearrangements with stable repertoires of genes and transposable elements in an invasive forest-pathogenic fungus": Supplemantary Figures: FigureS8_Demene2021_ReSub.pdf

**Figure S8:** Visualization of the mapping of the Nanopore (ESM015) or Sangers (EP155) reads on the ESM015 and EP155 genome assemblies around the major detected synteny breaks

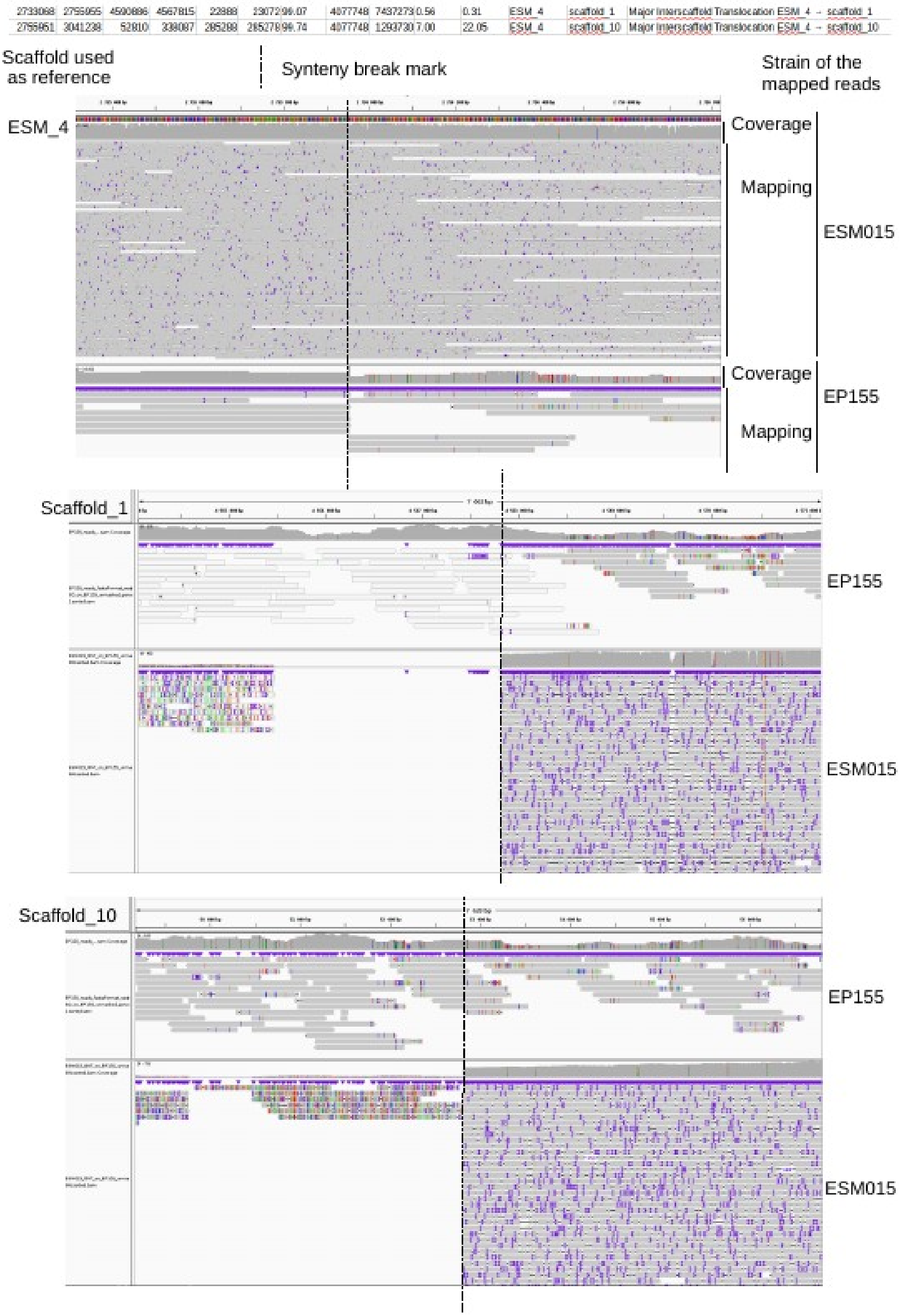

**Figure S8** (continued)

| | | | | | | | | | | | | | |
|---|---|---|---|---|---|---|---|---|---|---|---|---|---|
| 2412868 | 2511629 | 884193 | 785452 | 98762 | 98742 | 99.86 | 5381768 | 3395974 | 1.84 | 2.91 | ESM_2 | scaffold_7 | Major Interscaffold Translocation ESM_2 → scaffold_7 |
| 2511626 | 2522534 | 2543830 | 2554729 | 10909 | 10900 | 99.86 | 5381768 | 4337670 | 0.20 | 0.25 | ESM_2 | scaffold_5 | Major Interscaffold Translocation ESM_2 → scaffold_5 |

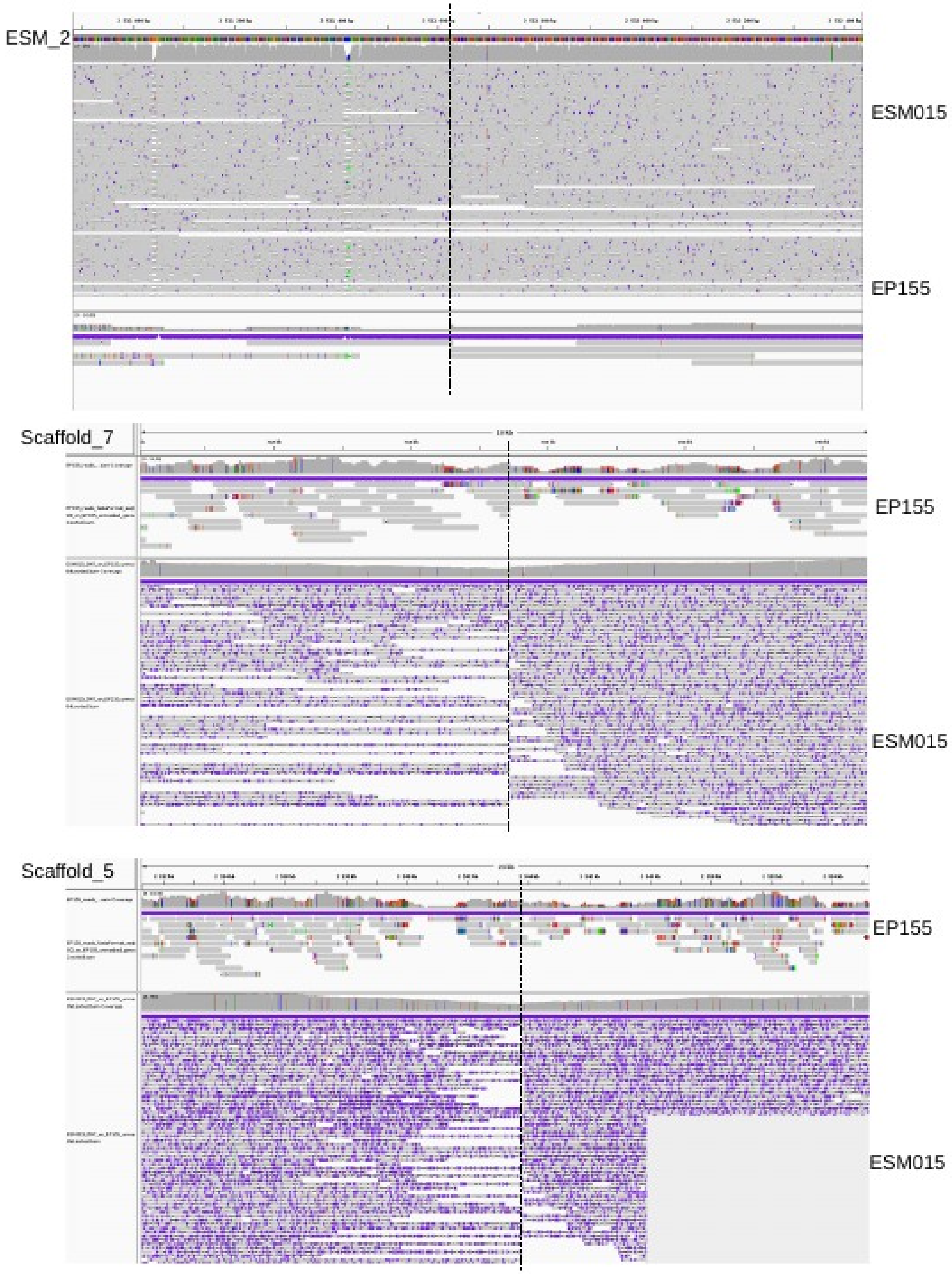

Figure S8 (continued)

| | | | | | | | | | | | | | |
|---|---|---|---|---|---|---|---|---|---|---|---|---|---|
| 3240299 | 3316747 | 3266023 | 3342486 | 76449 | 76464 | 99.80 | 5381768 | 4337670 | 1.42 | 1.76 | ESM_2 | scaffold_5 | Major Interscaffold Translocation ESM_2 → scaffold_5 |
| 3316746 | 3830593 | 3473810 | 3987657 | 513848 | 513848 | 99.83 | 5381768 | 5527719 | 9.55 | 9.30 | ESM_2 | scaffold_2 | Major Interscaffold Translocation ESM_2 → scaffold_2 |

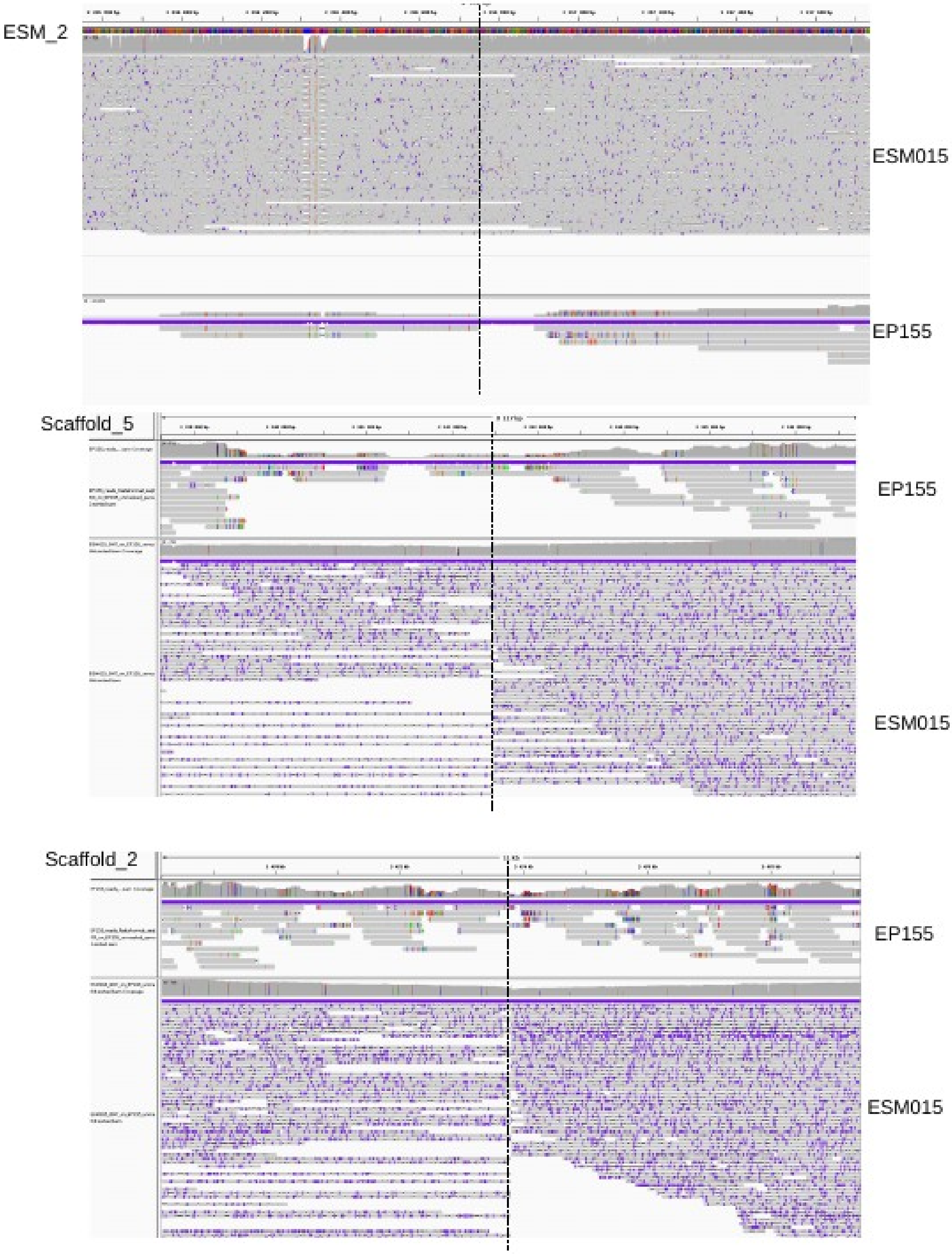

Figure S8 (continued)

| | | | | | | | | | | | | | |
|---|---|---|---|---|---|---|---|---|---|---|---|---|---|
| 1032112 | 1043922 | 3354296 | 3342486 | 11811 | 11811 | 99.99 | 2891089 | 4337670 | 0.41 | 0.27 | ESM_7 | scaffold_5 | Major Interscaffold Translocation ESM_7 → scaffold_5 |
| 1043921 | 1118430 | 3473810 | 3399285 | 74510 | 74526 | 99.88 | 2891089 | 5527719 | 2.58 | 1.35 | ESM_7 | scaffold_2 | Major Interscaffold Translocation ESM_7 → scaffold_2 |

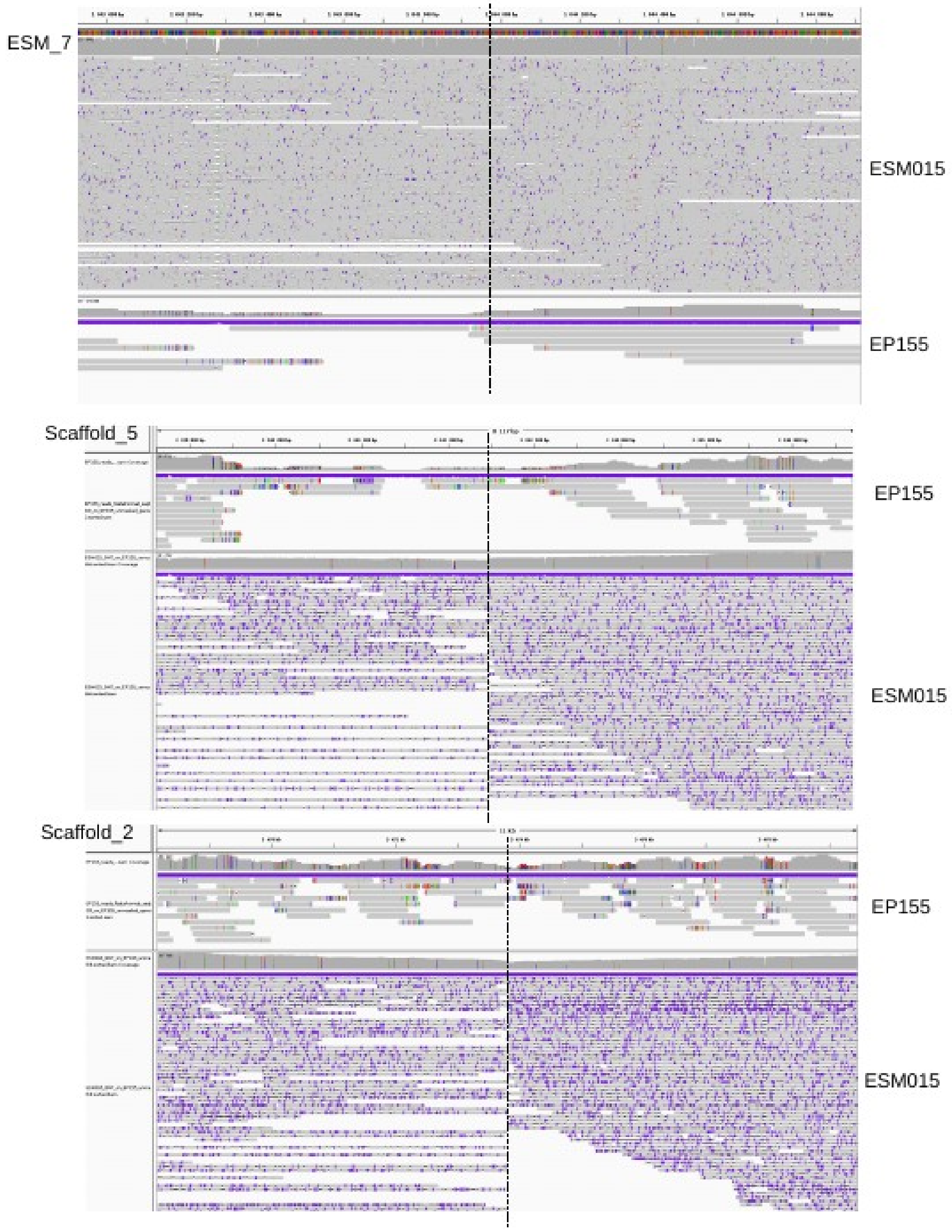

Figure S8 (continued)

| | | | | | | | | | | | | | |
|---|---|---|---|---|---|---|---|---|---|---|---|---|---|
| 1004321 | 1053110 | 736632 | 785455 | 48790 | 48824 | 99.89 | 3530110 | 3395974 | 1.38 | 1.44 | ESM_5 | scaffold_7 | Major Interscaffold Translocation ESM_5 → scaffold_7 |
| 1053107 | 1097962 | 2543833 | 2498955 | 44856 | 44879 | 99.90 | 3530110 | 4337670 | 1.27 | 1.03 | ESM_5 | scaffold_5 | Major Interscaffold Translocation ESM_5 → scaffold_5 |

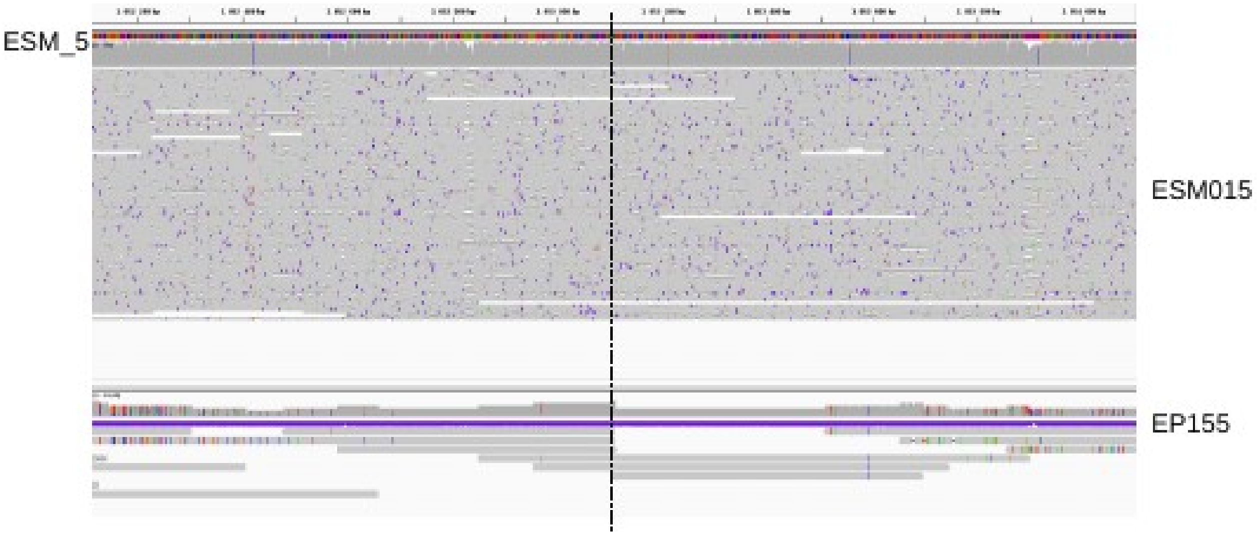


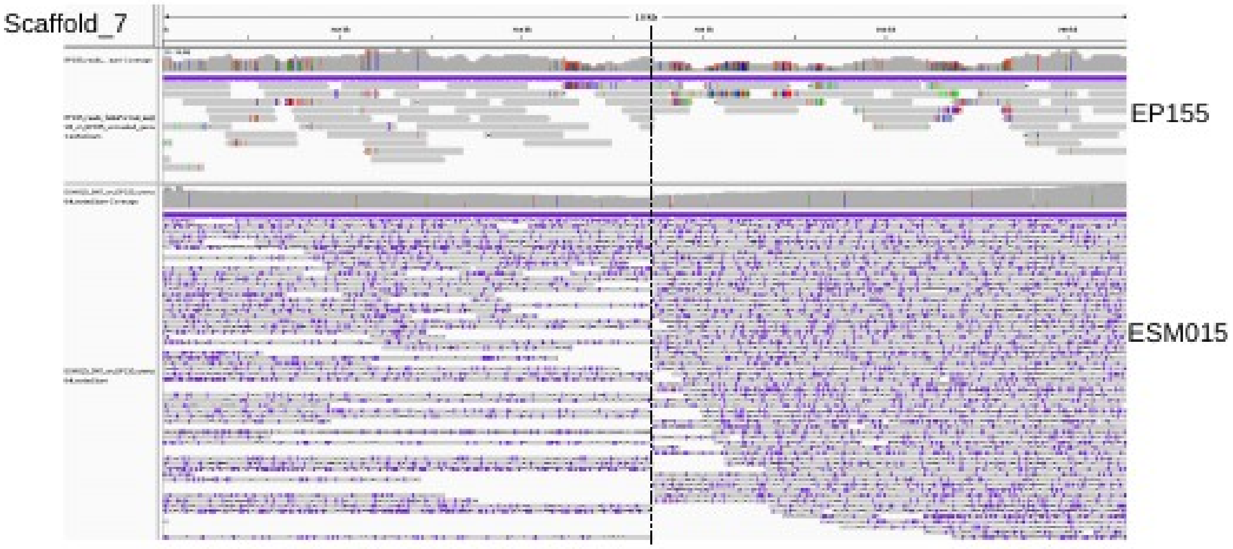


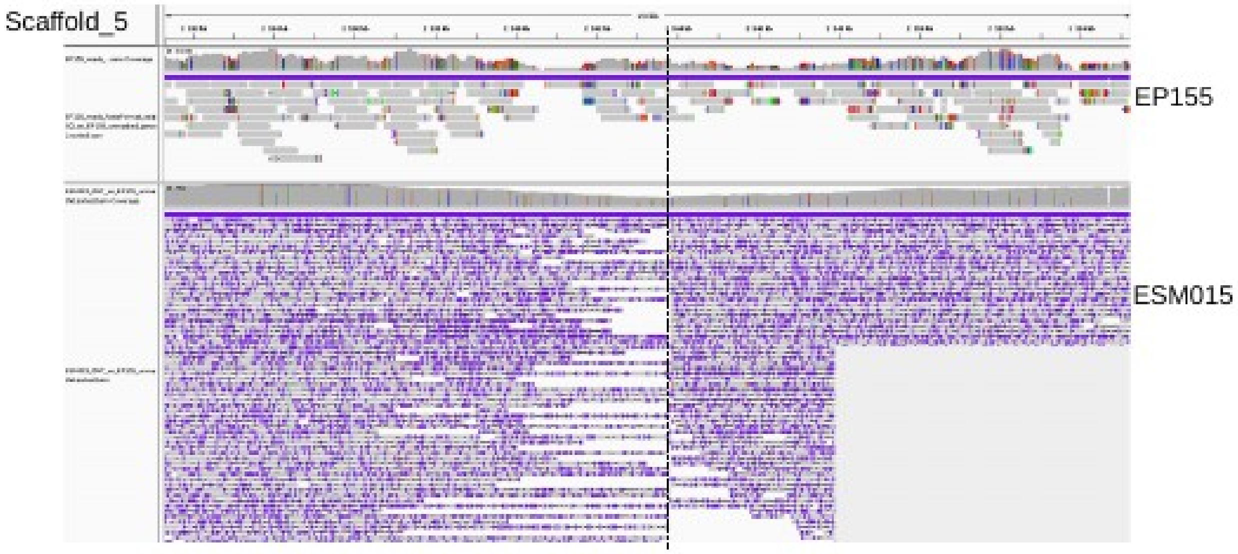

Figure S8 (continued)

| | | | | | | | | | | | | | |
|---|---|---|---|---|---|---|---|---|---|---|---|---|---|
| 1543121 | 1775927 | 2383767 | 2150857 | 232807 | 232911 | 99.87 | 3437609 | 3950794 | 6.77 | 5.90 | ESM_6 | scaffold_6 | Major intrascaffold translocation ESM_6 → scaffold_6 |
| 1775920 | 1883961 | 3350 | 111362 | 108042 | 108013 | 99.92 | 3437609 | 3950794 | 3.14 | 2.73 | ESM_6 | scaffold_6 | Major intrascaffold translocation ESM_6 → scaffold_6 |

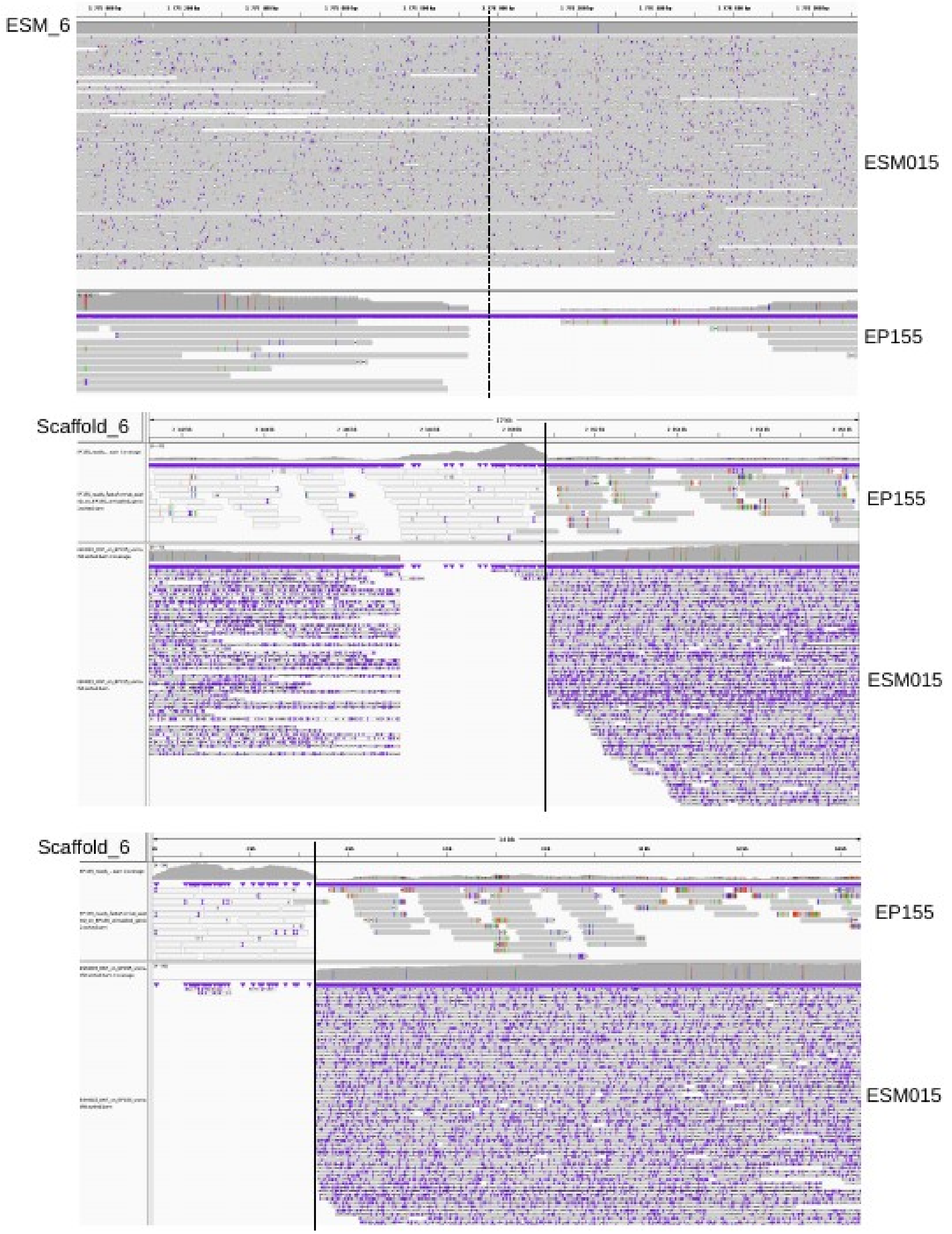

Figure S8 (continued)

| | | | | | | | | | | | | |
|---|---|---|---|---|---|---|---|---|---|---|---|---|
| 633307 | 682324 | 2090062 | 2139095 | 49018 | 49034 | 99.85 | 1663391 | 3950794 | 2.95 | 1.24 | ESM_10 | scaffold_6 | Major Interscaffold Translocation ESM_10 → scaffold_6 |
| 681854 | 690323 | 963434 | 974962 | 8470 | 8473 | 99.89 | 1663391 | 1008592 | 0.51 | 0.84 | ESM_10 | scaffold_11 | Major Interscaffold Translocation ESM_10 → scaffold_11 |

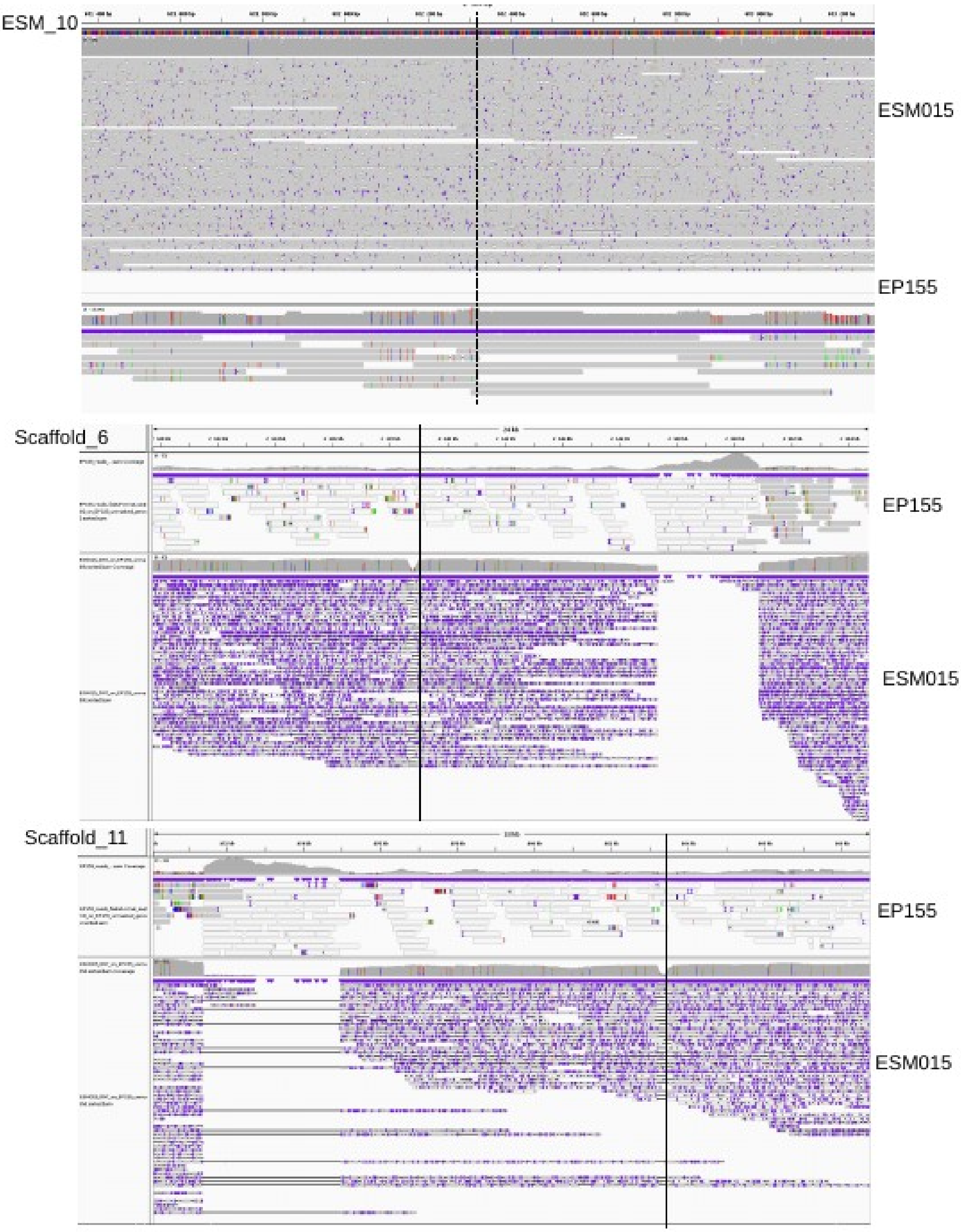

Figure S8 (continued)

| | | | | | | | | | | | | | |
|---|---|---|---|---|---|---|---|---|---|---|---|---|---|
| 75387 | 93013 | 1423790 | 1406164 | 17627 | 17627 | 97.48 | 599387 | 5527719 | 2.94 | 0.32 | ESM_11 | scaffold_2 | Major Intrascaffold Translocation ESM_11 → scaffold_2 |
| 115383 | 179603 | 591138 | 655400 | 64221 | 64263 | 98.97 | 599387 | 5527719 | 10.71 | 1.16 | ESM_11 | scaffold_2 | Major Intrascaffold Translocation ESM_11 → scaffold_2 |

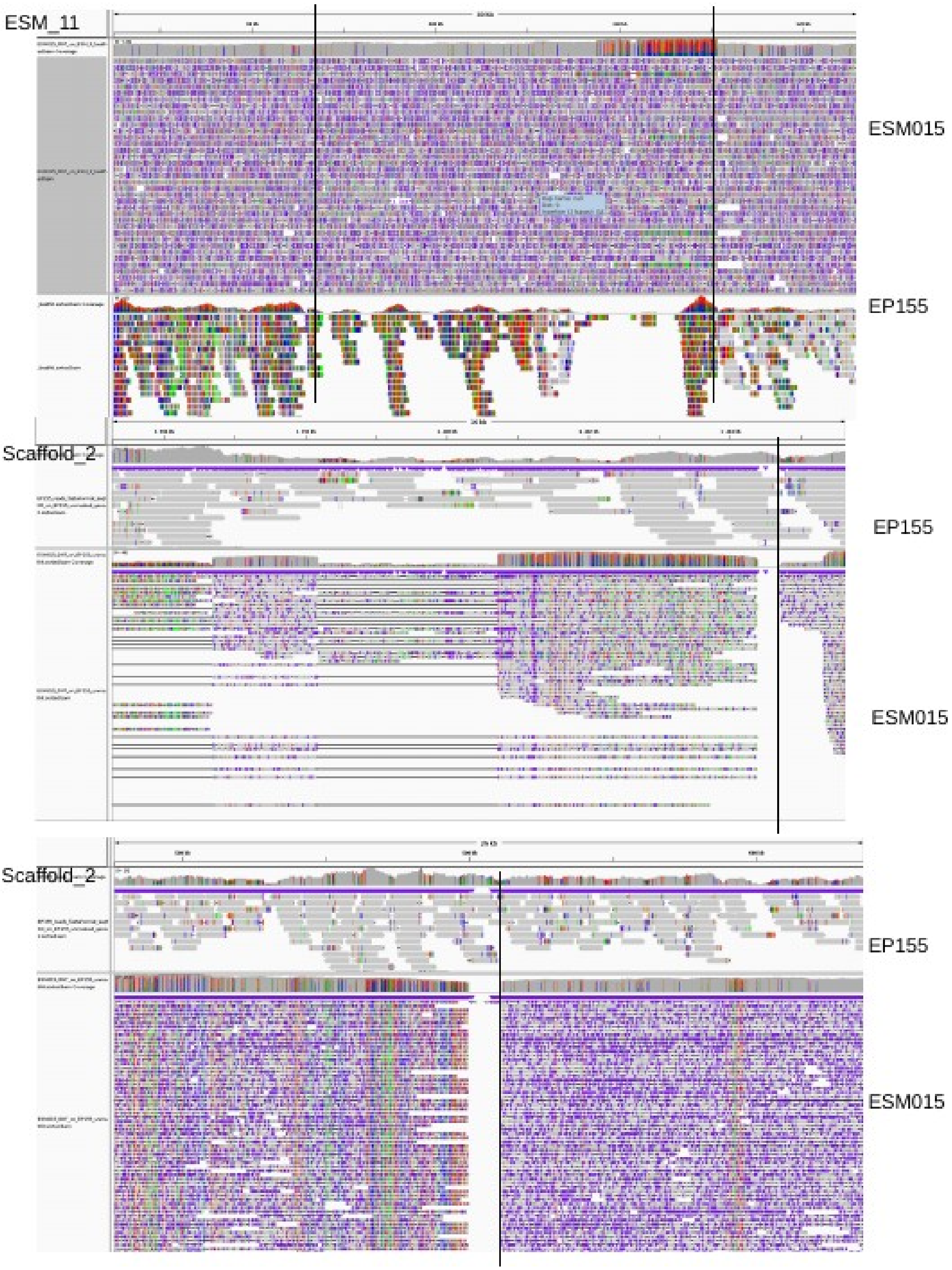

Figure S8 (continued)

| | | | | | | | | | | | | | |
|---|---|---|---|---|---|---|---|---|---|---|---|---|---|
| 187183 | 202667 | 574535 | 590007 | 15485 | 15473 | 97.07 | 500935 | 5527719 | 3.09 | 0.28 | ESM_12 | scaffold_2 | Major Intrascaffold Translocation ESM_12 → scaffold_[illegible] |
| 211902 | 234003 | 1377750 | 1355648 | 22102 | 22103 | 99.81 | 500935 | 5527719 | 4.41 | 0.40 | ESM_12 | scaffold_2 | Major Intrascaffold Translocation ESM_12 → scaffold_[illegible] |

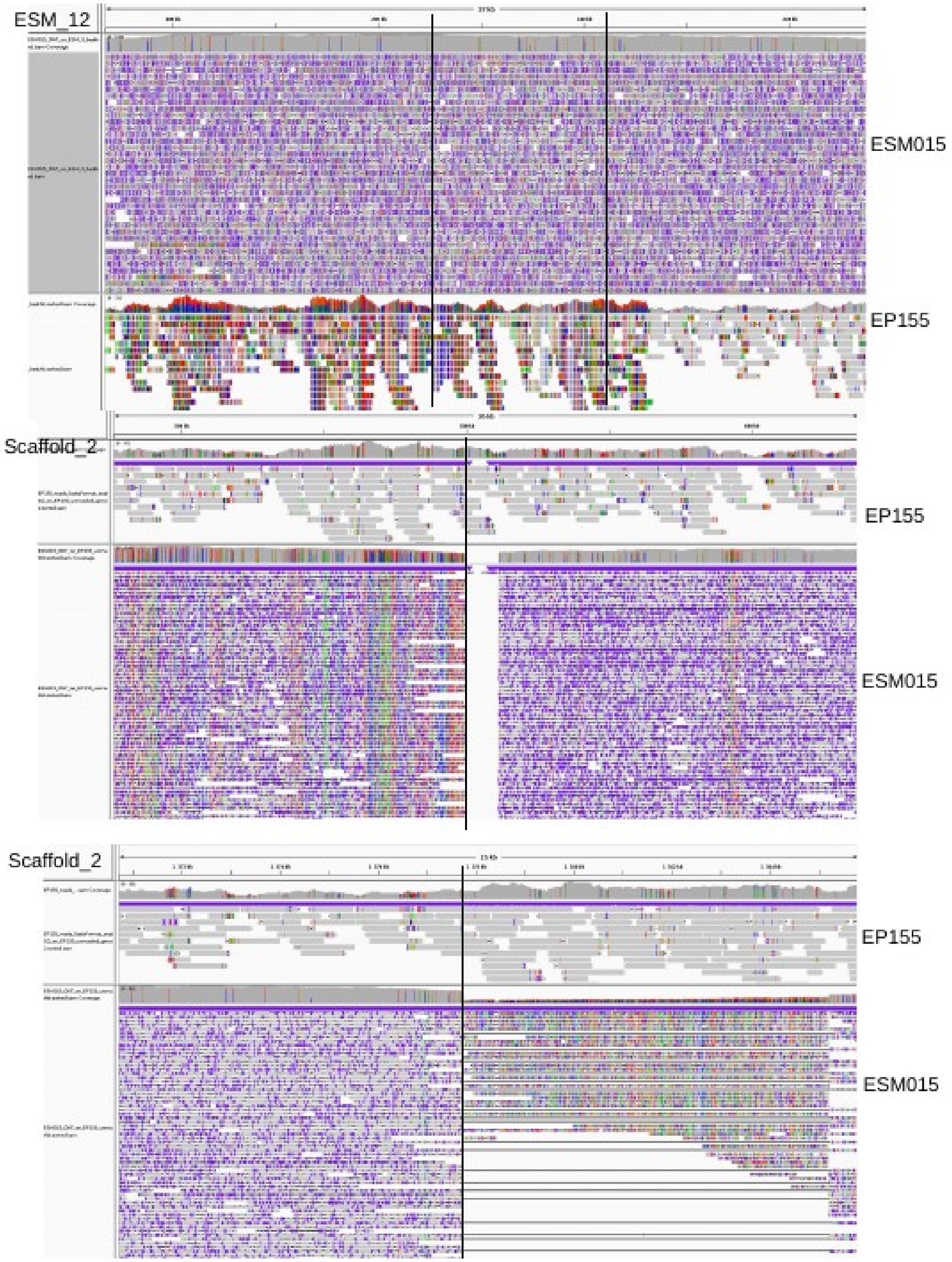
