## Supplementary figures and images for "Chromosomal rearrangements with stable repertoires of genes and transposable elements in an invasive forest-pathogenic fungus"

### FigureS6_Demene2021_ReSub.pdf

**Figure S6 :** estimated GC content in the ESM015 genome assembly

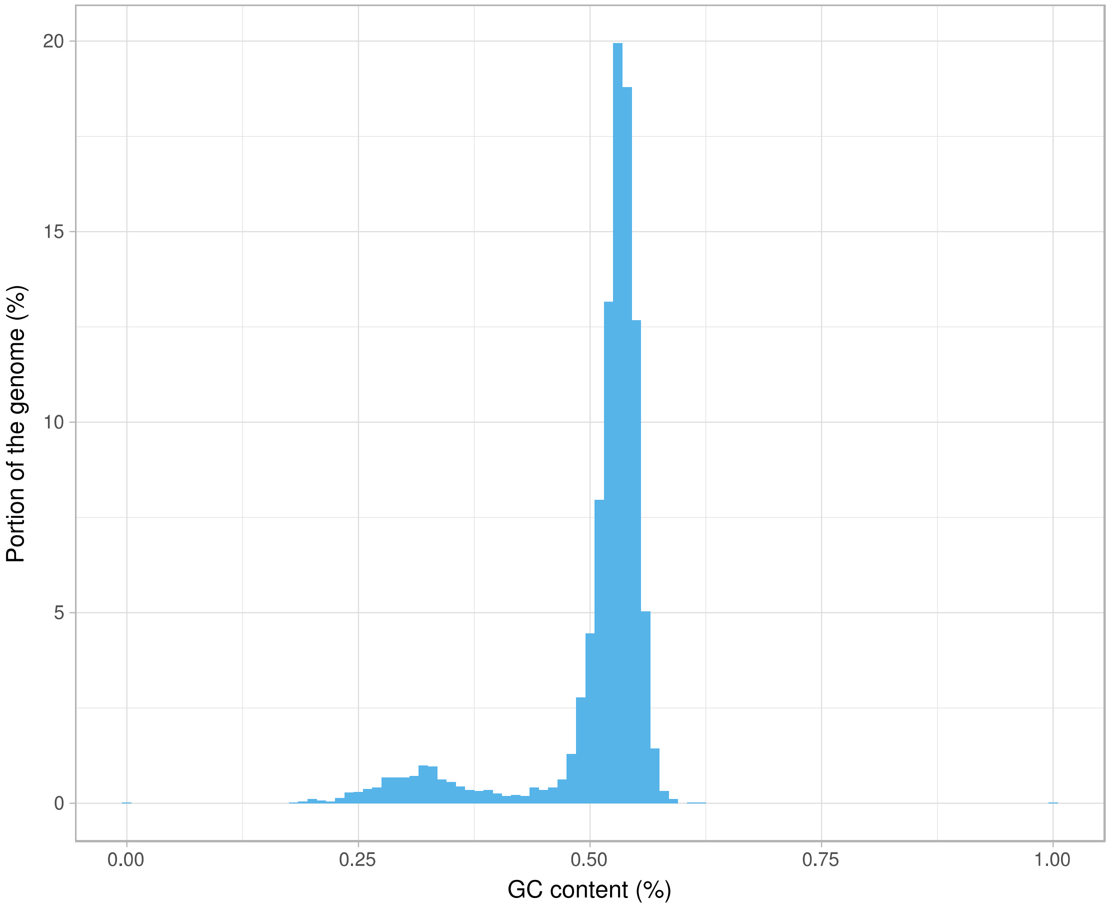
