## Supplementary material for "Chromosomal rearrangements with stable repertoires of genes and transposable elements in an invasive forest-pathogenic fungus": Supplemantary Material: SupMat3_Demene2021_ReSub.pdf

[Summary Statistics](#)

[Plots](#)

- Histogram of read lengths
- Histogram of read lengths after log transformation
- Weighted Histogram of read lengths
- Weighted Histogram of read lengths after log transformation
- Yield by length
- Read lengths vs Average read quality plot using dots
- Read lengths vs Average read quality plot using a kernel density estimation

### NanoPlot report

#### Summary statistics

| | |
|---|---|
| **General summary:** | |
| Mean read length: | 9399.2 |
| Mean read quality: | 8.0 |
| Median read length: | 8724.0 |
| Median read quality: | 8.9 |
| Number of reads: | 453365 |
| Read length N50: | 15460 |
| Total bases: | 4261267600 |
| **Number, percentage and megabases of reads above quality cutoffs** | |
| >Q5: | 387698 (85.5%) 4035.4Mb |
| >Q7: | 348625 (76.9%) 3725.5Mb |
| >Q10: | 57310 (12.6%) 762.7Mb |
| >Q12: | 4 (0.0%) 0.0Mb |
| >Q15: | 0 (0.0%) 0.0Mb |
| **Top 5 highest mean basecall quality scores and their read lengths** | |
| 1: | 12.4 (1106) |
| 2: | 12.3 (1884) |
| 3: | 12.2 (1394) |
| 4: | 12.1 (970) |
| 5: | 12.0 (2543) |
| **Top 5 longest reads and their mean basecall quality score** | |
| 1: | 80514 (6.8) |
| 2: | 79044 (2.8) |
| 3: | 77977 (6.8) |
| 4: | 76127 (6.6) |
| 5: | 75360 (7.3) |

#### Plots

##### Histogram of read lengths

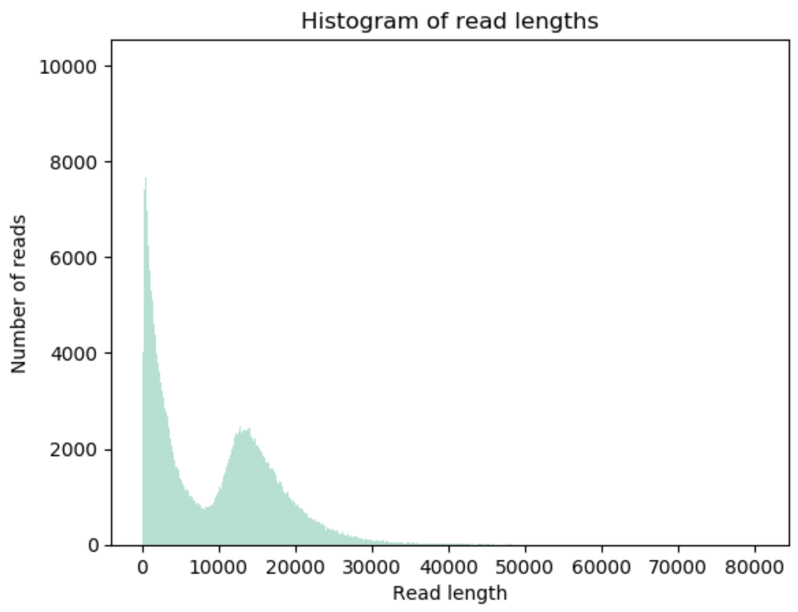


**Histogram of read lengths after log transformation**

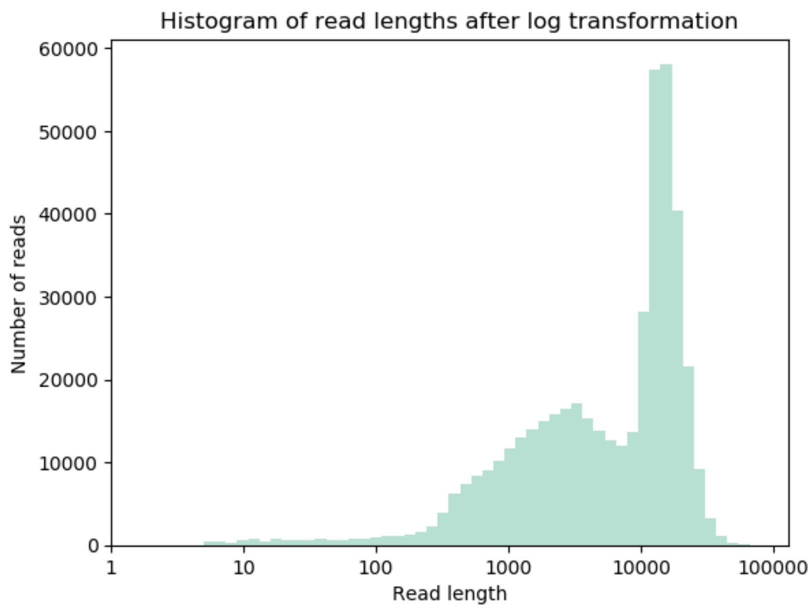


**Weighted Histogram of read lengths**

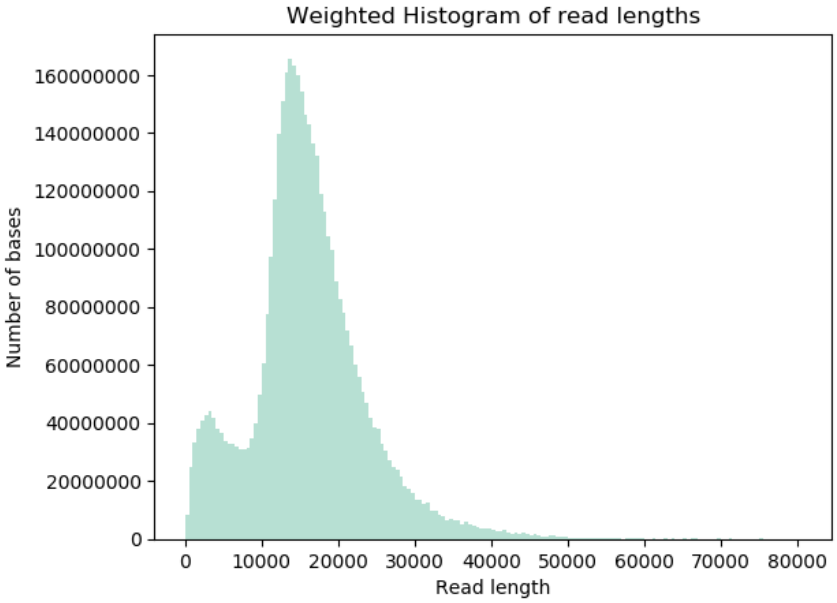


**Weighted Histogram of read lengths after log transformation**

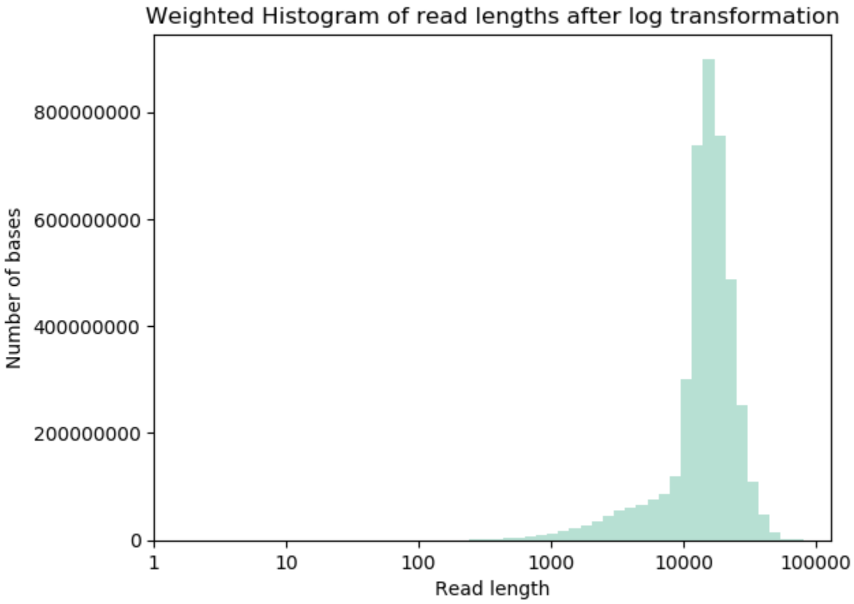


**Yield by length**

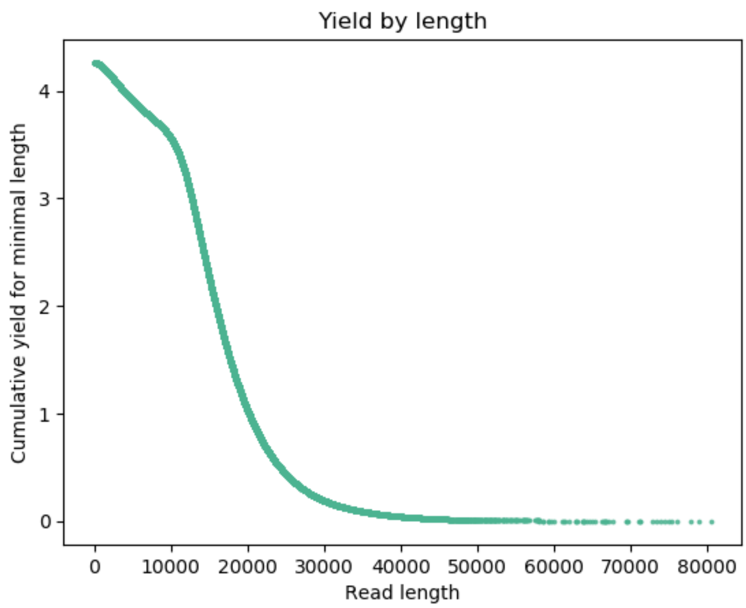


**Read lengths vs Average read quality plot using dots**

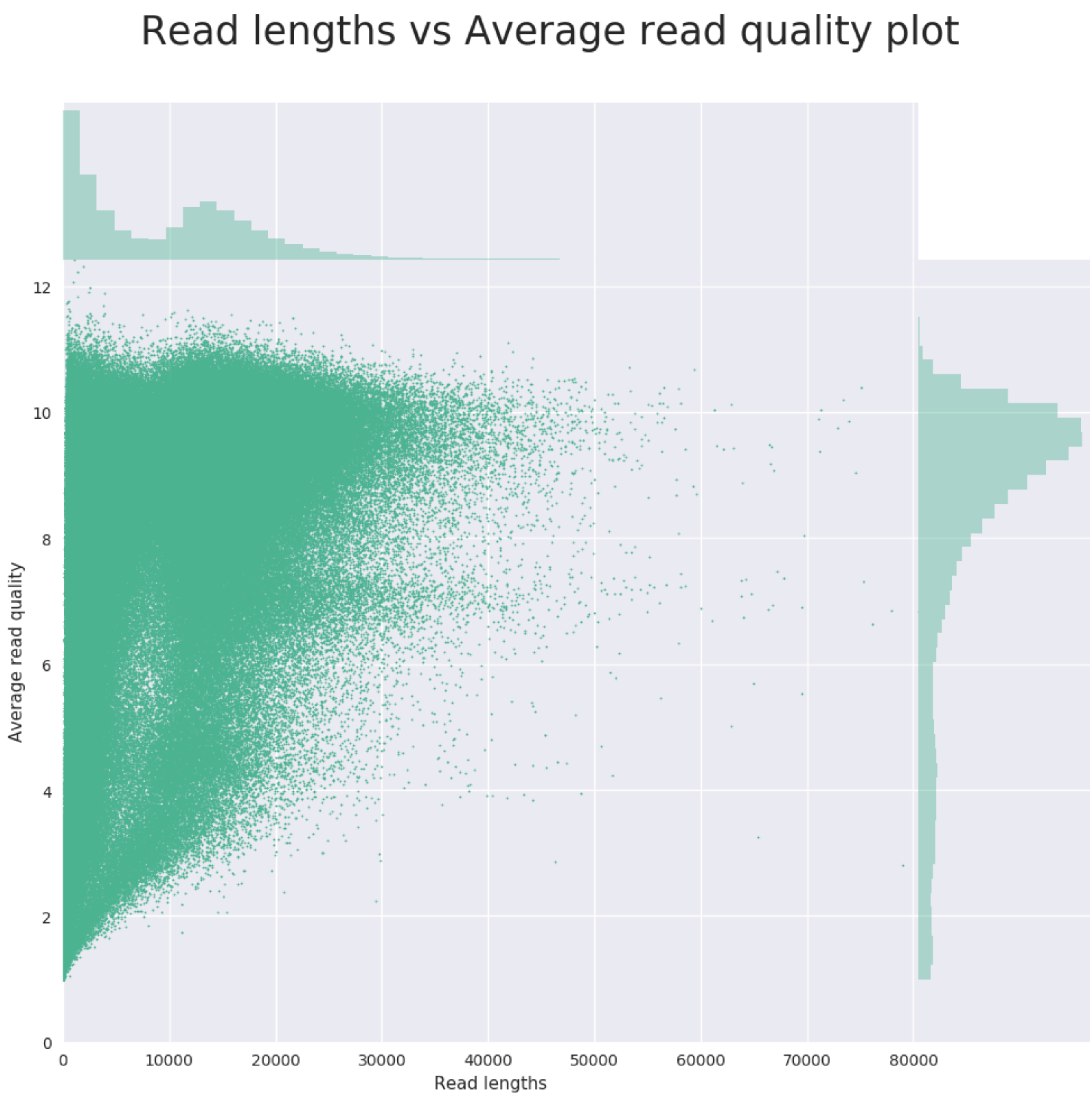


**Read lengths vs Average read quality plot using a kernel density estimation**

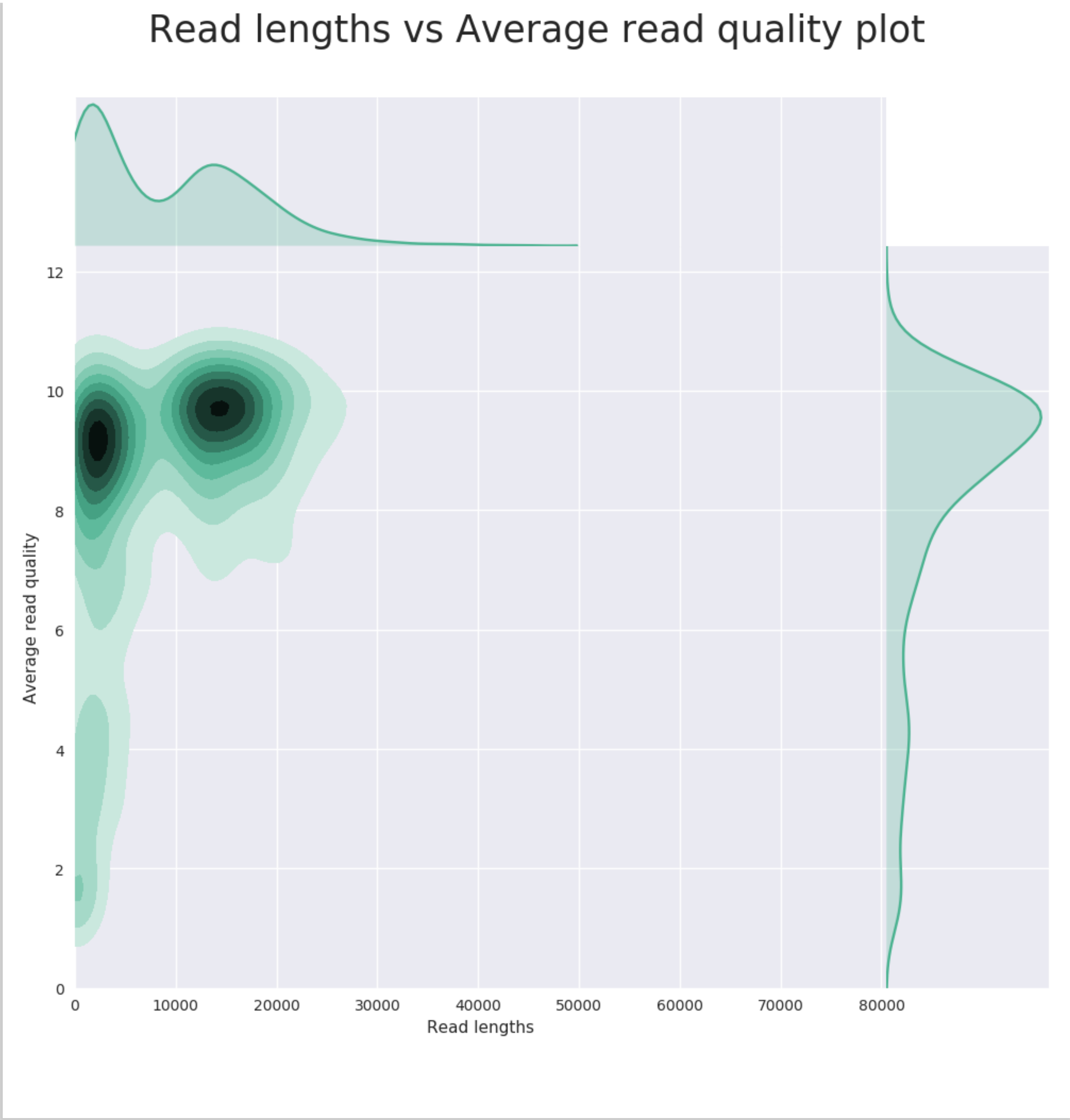
Read lengths vs Average read quality plot
Average read quality
Read lengths
0
2
4
6
8
10
12
10000
20000
30000
40000
50000
60000
70000
80000
